# Imageomics quantifies epidermal aging of sun-protected human skin and its rejuvenation by xenografting

**DOI:** 10.64898/2026.04.29.721704

**Authors:** Austin E. Y. T. Lefebvre, Ying Zheng, Ruifeng Yang, Fiona Lan, Arben Nace, Emilie Katz, Sergiy Libert, Cynthia Kenyon, Katie Podshivalova, George Cotsarelis

## Abstract

Rejuvenating aging human skin is a major therapeutic goal, but objective, quantitative measures of intrinsic aging are limited. We performed a cross-sectional histological study of UV-protected buttock and abdominal skin in Caucasian females aged 20 to 81 years to identify features that reliably track age. Epidermal thickness measured between rete ridges was unchanged, but rete ridge size declined linearly with age: ridges became shorter and thinner in both sites, though ridge number decreased only in the abdomen. Consistent with these structural changes, expression of integrin β4 (ITGB4), a putative stem-cell marker, was reduced in aged skin. We combined these biomarkers into a predictive model that estimated skin age more accurately than any single marker. To test whether the model detects longitudinal change, we analyzed aged abdominal skin before and after xenografting onto young or aged mice, a procedure previously reported to rejuvenate human skin in young but not aged recipient mice. Both the individual biomarkers and the model indicated rejuvenation regardless of host age; however, engraftment efficiency was lower in aged hosts, with surviving grafts showing younger histological phenotypes. These results provide quantitative criteria for assessing intrinsic skin aging and suggest that the process of engraftment itself induces rejuvenation-like changes.

## Introduction

The appearance of the skin is perhaps the most overt indicator of one’s age. Aging of the skin has been divided into “intrinsic” and “extrinsic”. Extrinsic is thought to be caused primarily by UV light whereas intrinsic is due to innate or endogenous biological processes, leading to recognized histologic and cellular changes in the skin (Quan 2023; Zhang and Duan 2018). In the aged, protected epidermis, the most pronounced histologic change is flattening of the epidermis from loss of rete ridges often referred to as “epidermal thinning” (Costello et al. 2023; Lavker et al. 1987; Quan 2023; Russell-Goldman and Murphy 2020).

The proportion of Ki-67 positive basal keratinocytes declines with age, presumably reflecting a loss of epidermal stem cells (Costello et al. 2023; Giangreco et al. 2008; Grove and Kligman 1983). Diminished proliferative capacity of basal keratinocytes is thought to contribute to “epidermal thinning” and age-related loss of rete ridges (Giangreco et al. 2008; Rübe et al. 2021; Zou et al. 2021). Chronologic aging correlates not only with reduction in the number of rete ridges but also manifests in discernible structural changes in residual rete ridges, which become shorter and thinner (Costello et al. 2023; Lavker et al. 1987). As the number of rete ridges diminishes, the dermal-epidermal junction flattens with age, and aged skin exhibits reduced expression of the epidermal adhesion molecule ITGB4 (Balin 1989; Langton et al. 2016; Lavker et al. 1987; Lavker 1979). Together, these histologic and molecular changes are indicative of age-associated remodeling of the dermal-epidermal junction, and compromised attachment between the epidermal and dermal compartments.

The dermis also expresses several intrinsic age-related changes. Pruning of the microvasculature of the dermis has been reported to occur with age (Balin 1989; Gunin et al. 2014), as indicated by reduced histological staining for various endothelial cell markers, including CD31 (Arnal-Forné et al. 2024; Gunin et al. 2014). This reduced capillary density presumably affects nutrient and oxygen delivery, as regression of vascular loops in the dermal papillae limits available surface area for exchange (Gilchrest 1982). Intrinsic aging also results in degradation and loss of collagen and elastin fibers, with associated remodeling and disorganization of the dermal extracellular matrix (ECM) (Marcos-Garcés et al. 2014; Mora Huertas et al. 2016; Shin et al. 2019; Varani et al. 2006). Changes in the macromolecular structure of the remaining fibers culminate in an increase in dermal stiffness (He et al. 2023; Jiang et al. 2025; Roig-Rosello et al. 2024).

The study of intrinsic aging in skin is complicated by several factors, including UV exposure and differences across anatomical sites. UV radiation arrests proliferation in the interfollicular epidermal stem cell population (Panich et al. 2016) and upregulates expression of matrix metalloproteinases which degrade collagen, elastin, laminin, and other hemidesmosome proteins that anchor the dermal-epidermal junction (Doleckova et al. 2024; Feng et al. 2024). In addition, the morphology of human skin differs across anatomic sites, so conclusions drawn from single-site studies are not necessarily generalizable (Balin 1989).

While any observed morphologic change or biomarker discussed above might individually allow for estimation of chronologic age, consideration of a combination of markers would be more comprehensive and might allow for a more accurate estimation of skin age and health. In addition, these age-dependent biomarkers have not yet been applied to evaluating longitudinal changes in skin integrity, particularly in response to conditions thought to induce rejuvenation or protection from disease states. Prior work has reported that, in a xenograft model, aged human skin grafted onto young immunodeficient host mice exhibits signs of rejuvenation with respect to a series of key biomarkers associated with intrinsic skin aging (Keren et al. 2022). In demonstrating that rejuvenation of aged human skin is possible, this work raises the additional question of how to quantify intrinsic skin aging – an inherently complex and variable phenotype. Such quantification would permit not only a systematic characterization of skin age, but also a standardized means of evaluating the skin’s response to rejuvenation therapies.

Despite the general agreement on the major histologic hallmarks of intrinsic skin aging, most studies have operationalized these phenotypes through manual measurements and semi-quantitative scoring applied to a limited number of representative fields of view, an approach that is time-intensive, can be subjective, and is vulnerable to both inter-observer variability and bias introduced by field selection in a spatially heterogeneous tissue such as skin (Balin and Pratt 1989; Bencze et al. 2021; Brown 2017; Knoblaugh and Himmel 2019). More recently, whole-slide imaging and digital pathology have made it practical to interrogate substantially larger tissue areas on routine H&E sections, enabling standardized, reproducible morphometric quantification (e.g., epidermal thickness distributions, rete ridge density and geometry, and compartmental area fractions) rather than relying on small, manually selected regions (Aeffner et al. 2019; Kumar et al. 2020). In skin specifically, digital approaches can yield more reliable and accurate measurements of dermal and epidermal thickness than traditional methods, supporting a shift toward quantitative readouts as primary endpoints (Turin et al. 2018). Parallel advances in open, scalable software ecosystems for whole-slide analysis (e.g., QuPath) have lowered the barrier to batch processing and objective feature extraction across cohorts, helping to standardize analyses and reduce dependence on manual workflows (Bankhead et al. 2017). Together, these developments suggest that integrating broad-area, image-derived morphometrics from H&E with established aging-associated biomarkers could provide a more comprehensive and less biased framework for estimating “skin age” and for measuring longitudinal change, but baseline distributions across defined age bands in UV-protected, anatomically consistent human skin-and principled methods to combine multiple markers into a predictive model-remain incompletely established.

In this study, we sought to undertake an unbiased identification of age-related changes in histological skin features coupled with a rigorous quantification of established biomarkers of aging in UV-protected skin from two different anatomical sites with the goal of establishing a baseline for each biomarker across designated age bands, and then integrating these individual biomarker parameters into a novel, predictive model of skin age that might more accurately characterize skin health and aging. We then tested this novel model in a previously described xenograft system in order to assess with greater precision the extent of skin rejuvenation. Surprisingly, we found that aged human skin appeared rejuvenated after grafting onto young or old mice.

## Results

### Morphological epidermal changes with age

Epidermal features were extracted from digitized H&E-stained skin sections sampled from the buttock (young, 20-30 years; middle-aged, 40-50; old, 60 and over) and abdomen (ages 20-30 and 70+; Supplementary Figure S1, Supplementary Table S1, Supplementary Table S2), all from Caucasian female individuals, using an image-processing pipeline that needs no manual annotation of these sections (Figure 1a-d). In brief, whole-slide RGB images were downsampled to identify tissue regions, which were segmented into individual slices based on connected components (Figure 1a). Each slice was processed at high resolution to isolate the epidermis using color filtering, optical density deconvolution, and morphological operations (Figure 1b,c). Epidermal regions were skeletonized and converted into graph representations to identify the epidermal midline and rete ridges (Figure 1d,e). Morphometric features-including thickness profiles, ridge lengths, and dilation factors-were extracted from each valid region and combined with manually extracted immunofluorescence (IF) features for quantitative analysis (Figure 1f-h). A detailed description of the image-processing methodology is described in the methods section. Linear regression models were independently fit for both buttock (Figure 2a,c) and abdomen (Figure 2b,d) samples to assess age-associated trends in morphometric features of the epidermis, specifically focusing on metrics derived from rete ridge geometry and epidermal thickness. All regression results, with slope confidence intervals, effect sizes and q-values within each modality and site, are given in Supplementary Table S3; associations with R² < 0.10 are reported but treated as weak.

**Figure 1.**
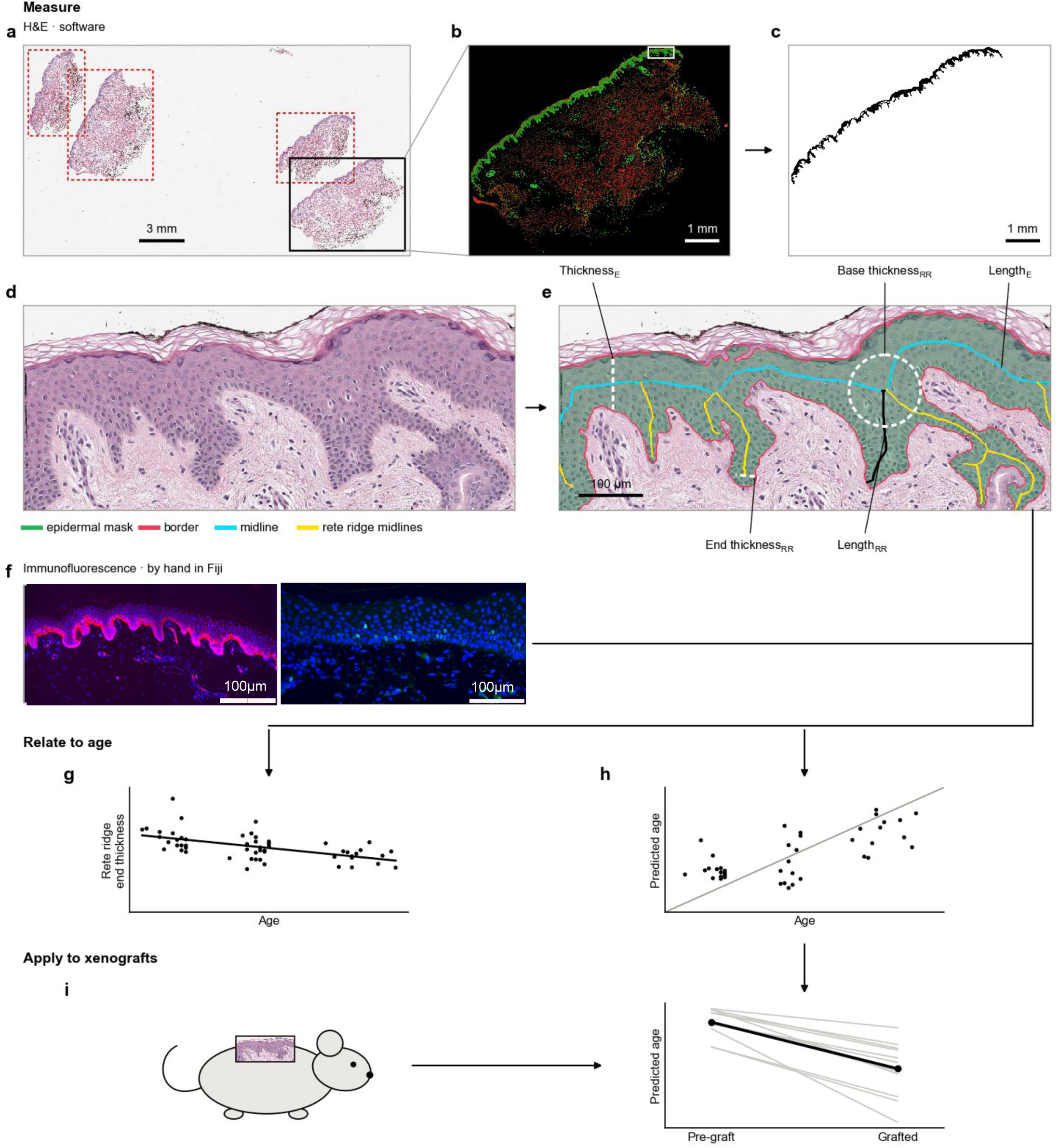
The measurement pipeline from stained section to predicted skin age. a. &E section segmented by software into 47 features per section b. optical density after stain deconvolution c. epidermal mask d. Boxed epidermis as scanned e. Segmentation of the epidermis with example epidermal (E) and rete ridge (RR) features (dashed white line): epidermal thickness, rete ridge base thickness (dashed white circle) and end thickness (dashed white line), with rete ridge length as the black path. f. ITGB4 intensity and Ki67-positive cells, measured by hand in Fiji. g. Rete ridge end thickness regressed on age. h. Age model predictions.

**Figure 2.**
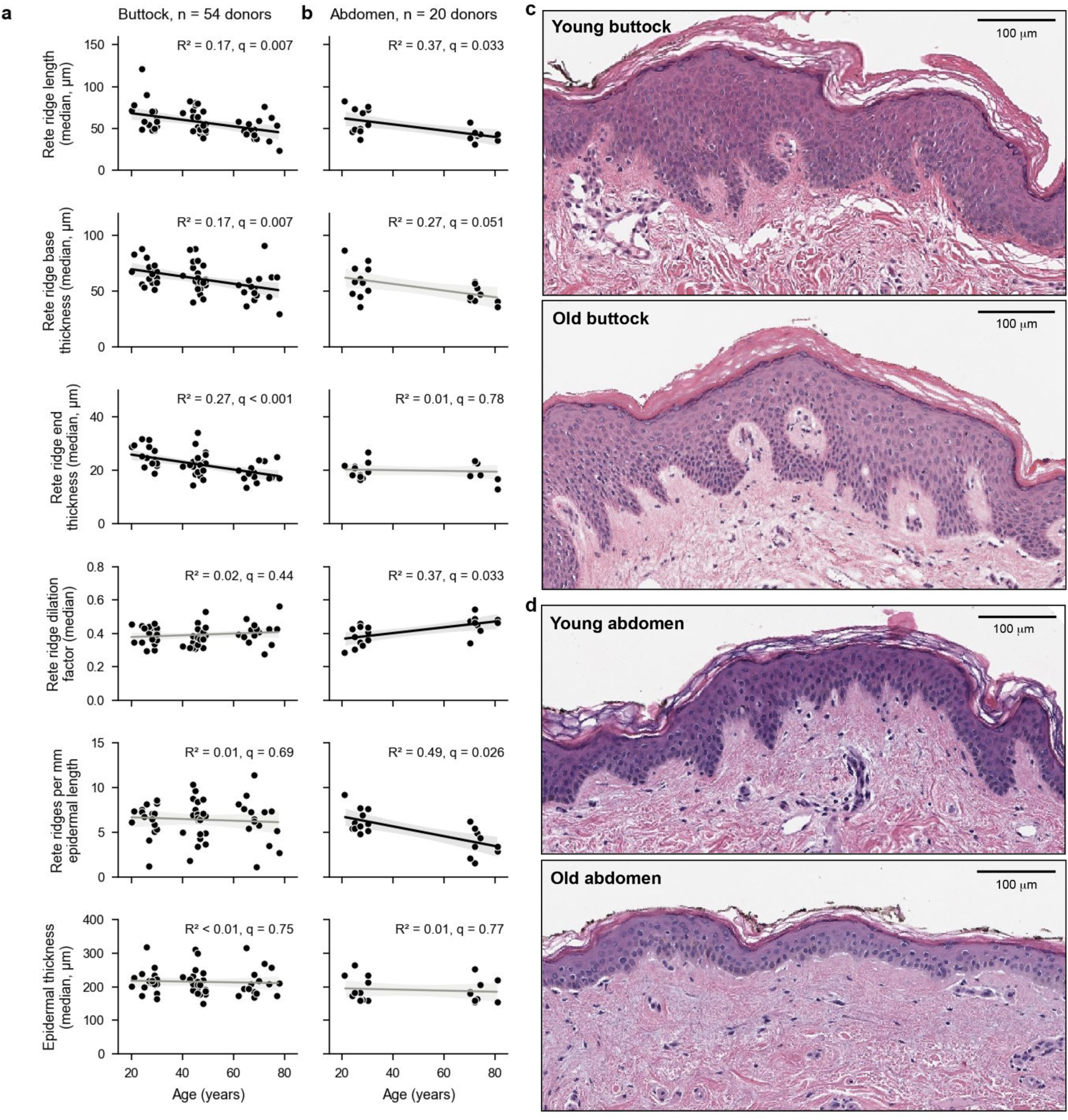
Rete ridge morphology changes with age. a, b. Epidermal morphometrics of buttock (a; n = 54 donors) and abdominal (b; n = 20 donors) skin against donor age. Each point is one donor, the biological replicate: the median over the tissue sections on its slide (2 to 8 per buttock donor, median 4; 1 to 8 per abdominal donor, median 2). Lines are least-squares fits with 95% confidence bands. Each panel gives R² and the Benjamini-Hochberg q across the site’s 47 H&E features; fits with q ≥ 0.05 are grey. c, d. Representative H&E images of buttock (c) and abdominal (d) skin from young (top) and old (bottom) donors. Scale bars: 100 µm.

Across both anatomical sites, no significant age-associated change was detected in inter-rete ridge epidermal thickness (Figure 2a and b; manual measurements of epidermal area and rete ridge number are shown in Supplementary Figure S2). Several morphological features of the rete ridges showed statistically significant linear relationships with age after correction for multiple testing within each site (Supplementary Table S3). Rete ridge length decreased with age at both sites (median length, q = 0.007 in the buttock and q = 0.03 in the abdomen). Rete ridge base thickness also declined, in the buttock (median, q = 0.007) and in the abdomen (mean, q = 0.04; median, q = 0.05). The interquartile range (IQR) of base thickness decreased in the buttock (q = 0.02), with the same trend in the abdomen (q = 0.05), indicating reduced heterogeneity in ridge base dimensions with increasing age. Rete ridge end thickness thinned with age in the buttock (median, q < 0.001) and not in the abdomen (q = 0.78). The ratio of rete ridge end thickness to base thickness (dilation factor) showed a different pattern. In the abdomen, the median dilation factor increased with age (q = 0.03), indicating a progressive flattening of rete ridges. These findings suggest that rete ridges become shorter and relatively less sharp with age in abdominal, but not buttock skin. Lastly, the number of rete ridges per unit epidermal length was the strongest negative age association in the abdomen (r = −0.70, q = 0.03), indicating a marked reduction in rete ridge density in aged abdominal skin. In contrast, surprisingly, rete ridge density remained constant with age in the buttock skin (q = 0.69).

Collectively, these results demonstrate that while overall epidermal thickness remains stable with age in UV-protected skin, rete ridge morphology-including length, thickness, and density-undergoes consistent and measurable degeneration, with the abdomen showing more pronounced and coherent age-associated trends than the buttock, highlighting the differential susceptibility of skin from different anatomical locations to age-related changes.

### Changes at the epidermal-dermal interface with age

To test whether age-related flattening of rete ridges is mirrored by changes in the proliferative basal layer, we quantified basal adhesion, proliferation, and papillary microvasculature in sun-protected buttock skin from the same age-stratified Caucasian female cohort analyzed above (Supplementary Table S1, Supplementary Table S2). We measured integrin beta-4 (ITGB4) intensity in the basal epidermis to index hemidesmosomal attachment, counted Ki67-positive nuclei to estimate basal proliferative output, quantified collagen 17 intensity, and segmented CD31-positive objects in the papillary dermis to capture microvascular architecture. Immunofluorescence sections were compartment-annotated, features were summarized for each slice, and association was tested with a separate linear regression of the donor-level measurement on chronological age. All IF-derived features are summarized in Supplementary Table S4.

#### ITGB4 Expression in the Epidermis Decreases with Age

Integrin β4 (ITGB4) is specifically expressed in the basal epidermis. It is a component of hemidesmosomes, playing a crucial role for attaching the epidermis to the basement membrane. To assess if ITGB4 expression diminishes with age, contributing to the weakened epidermal-dermal junction observed in aging skin, we quantified its expression in the epidermis across the age-stratified cohort (Figure 3a). A linear regression model was fit to the mean ITGB4 intensity values across age. The analysis revealed a statistically significant, negative association between age and ITGB4 expression (n = 52 donors, r = −0.62, R² = 0.38, q < 0.0001; Figure 3b; Supplementary Table S4).

**Figure 3.**
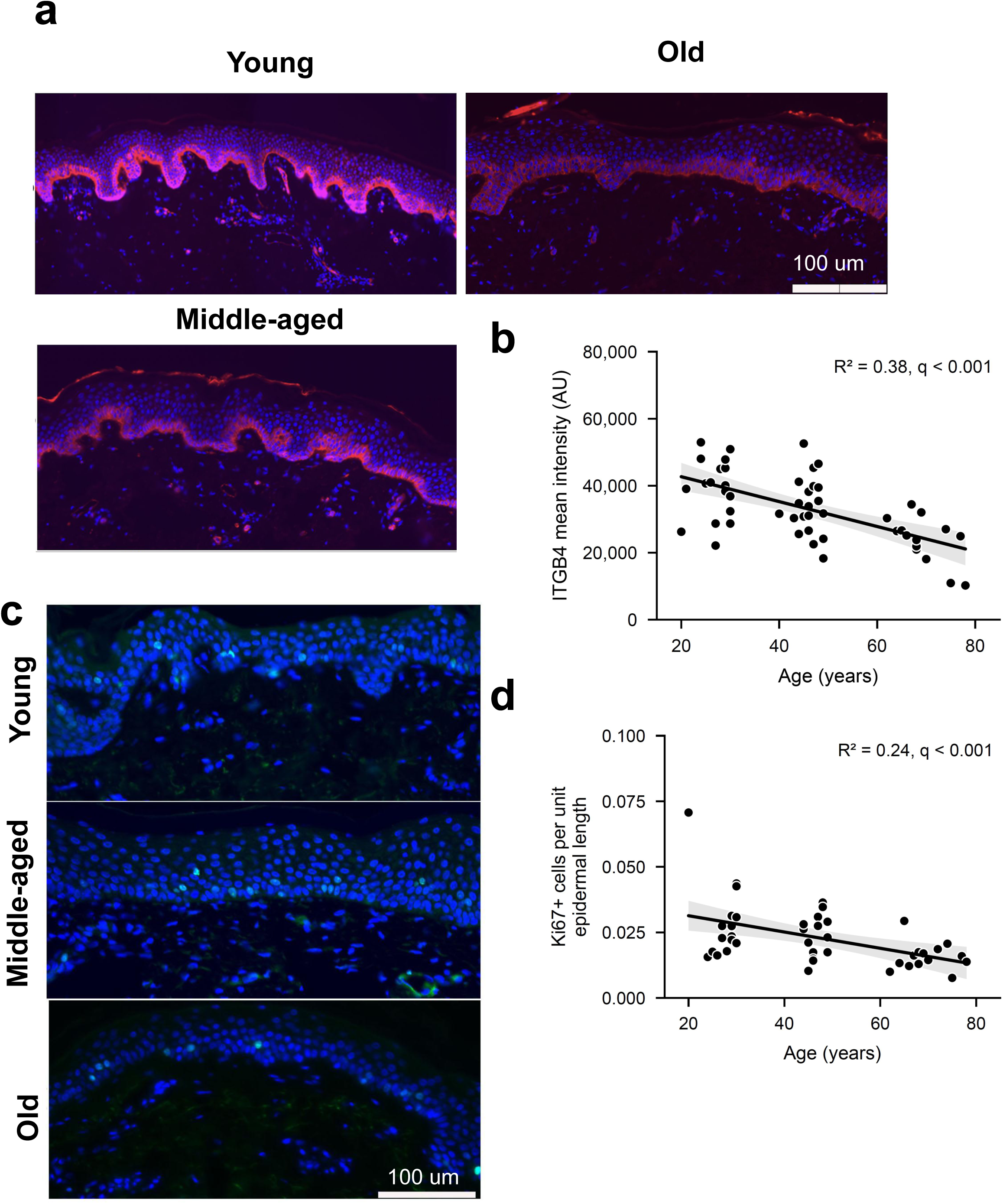
Age-related changes in the basal epidermal layer. a. ITGB4 immunostaining (pink), which marks the basement membrane zone, in buttock skin from young, middle-aged and old donors. b. ITGB4 mean intensity in the basal epidermis, measured in Fiji, against donor age (n = 52 donors). c. Ki67 immunostaining (green) in buttock skin from young, middle-aged and old donors. d. Ki67+ cells per unit epidermal length, counted by hand, against donor age (n = 43 donors). In b and d each point is one donor (one section each; no technical replicates); lines are least-squares fits with 95% confidence bands. R² and Benjamini-Hochberg q are shown; ITGB4 is tested alone (q equals p), Ki67 across its two measures. Counts exclude nine donors with a tissue-quality note. Scale bars: 100 um (a), 50 um (c).

#### Ki67 Proliferation in the Epidermis Decreases with Age

To evaluate age-related changes in epidermal proliferation, we counted Ki67+ cells within manually segmented epidermal regions from buttock skin sections across the age-stratified cohort (Figure 3c) and expressed the counts in two ways, per unit epidermal length and per rete ridge. Both declined with age: Ki67+ cells per unit epidermal length (n = 43, r = −0.49, R² = 0.24, p = 0.0009; Figure 3d) and Ki67+ cells per rete ridge (r = −0.50, R² = 0.25, p = 0.0006), and both associations survive correction across the two Ki67 measures (q = 0.0009). The association is weaker when the nine donors excluded from all other analyses for specimen-quality notes are included (per unit length: n = 51, R² = 0.086, p = 0.037, q = 0.063; Supplementary Table S4).

We also performed an exploratory analysis of the papillary dermal microvasculature by CD31 immunofluorescence (Supplementary Figure S3, Supplementary Figure S3, Supplementary Table S4). Mean vessel solidity rose and mean minor axis length fell with donor age at nominal significance (p = 0.045 and 0.044; n = 30 donors), but no CD31 feature survived correction across the 43 features (smallest q = 0.087), so these data do not establish an age-related change in the papillary microvasculature. Because CD31 analysis addresses a different tissue compartment (dermis vs. epidermis) than the rest of the study and cannot be extended to the xenografts, in which graft vasculature becomes chimeric with host murine endothelium (Demarchez et al. 1987; Young et al. 1996), we present it as a preliminary observation and do not include it in the age prediction model described below. We also found no significant changes in Col XVIIa with age (Supplementary Figure S5).

### Predictive Modeling of Skin Age

We assessed whether histomorphometric and basal molecular features predict chronological age in sun-protected buttock skin via a Random Forest regressor (Figure 4a). With feature selection and polynomial expansion repeated inside each cross-validation fold, the model predicted age with a mean absolute error (MAE) of 9.73 years (95% CI 9.42 to 10.03) and R² of 0.567 (0.536 to 0.599), better than any individual feature (Figure 4b).

**Figure 4.**
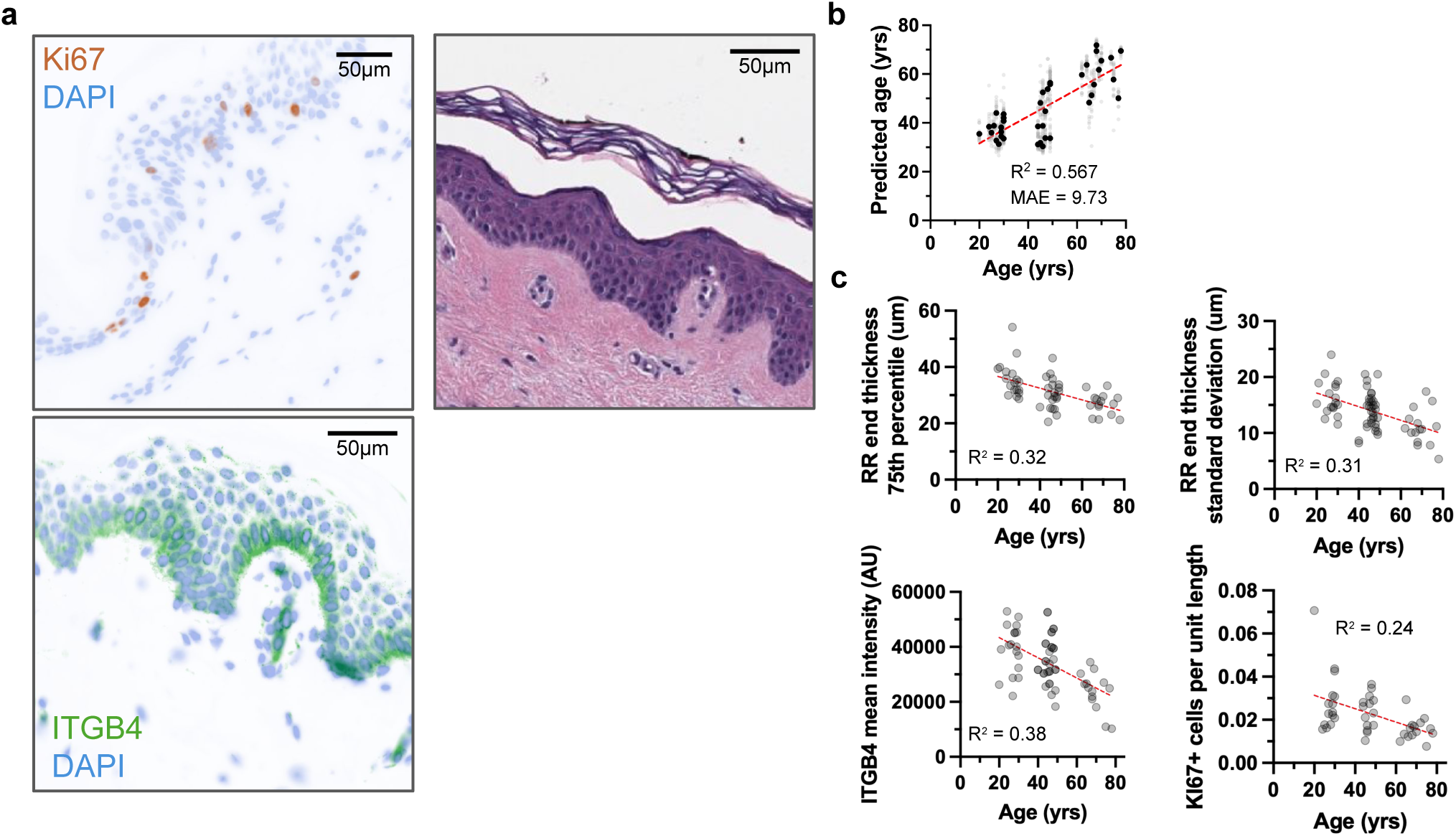
Skin age predictive modeling. a. Representative Ki67 and ITGB4 immunofluorescence and H&E staining. b. Predicted against actual age for the reduced polynomial random forest (n = 41 donors). Light grey points, each donor’s out-of-fold prediction in each of 20 repeats of 5-fold cross-validation, with feature selection and expansion inside each training fold; black points, per-donor means; dashed line, [identity line / least-squares fit]. MAE and R² are means over the 20 repeats (95% CI 9.42 to 10.03 years; 0.536 to 0.599); the per-donor means alone give 9.46 years and 0.614. c. Single model features against donor age; each point is one donor (n = 54, 54, 52, 43); R² from least-squares fits.

Feature importance concentrated on rete ridge end thickness (Supplementary Figure S6). The 75th percentile of end-thickness was the single most informative predictor, followed by its standard deviation and range; together these three metrics accounted for ∼40% of total importance. Additional contributors included ITGB4 mean intensity, Ki67-positive cell density, and other rete-related thickness metrics, particularly those describing end morphology (Figure 4c).

### Predictive modeling shows rejuvenation of human skin xenografted into mice

Next, we wanted to test whether the predictive model could be applied to quantify the effect of an anti-aging intervention. It has been previously reported that transplanting aged human skin onto young immunodeficient mice morphologically rejuvenates the xenograft, whereas transplanting the same skin onto old mice has no such effect (Keren et al. 2022).

#### Xenograft survival is impaired in old host mice

To evaluate whether the age of the host environment influences the epidermal age of the skin graft, for each of seven elderly female donors (66 to 75 years old, mean 70.0; Supplementary Table S5), adjacent samples from the same surgical specimen of abdominal skin were xenografted onto both young and old immunodeficient mice. Acute mortality trended higher when the surgery was performed on old animals (17%) as compared to young animals (3%), although the difference was not statistically significant (Figure 5a). Within the surviving animals, we examined the engraftment success rate. Some xenografts clearly failed to establish, as evidenced by gross morphology and H&E staining, showing presence of only mouse skin at the surgical site (Figure 5b).To distinguish definitively between human and mouse skin, we used species-specific lamin staining. In successful or partially successful xenografts (Supplementary Figure S7), only human cells were detected in the epidermis, while in failed xenografts (Supplementary Figure S8), no human cells were detected, including in the areas with thick epidermis, which could be mistaken for human skin by H&E staining alone. In most xenografts, we detected variable amounts of infiltration of mouse cells into the dermis of the xenografted skins (Figure 5c-d). Grafts in which areas of mouse skin adjoined the human skin (Figure 5c) were scored as partially successful, and grafts with only human skin at the site (Figure 5d) as fully successful. Overall, the proportion of failed xenografts was significantly higher in old animals (60%) than in young animals (29%; Figure 5e). For all further histological analyses, only partially or fully successful xenografts were included after excluding any ingrown mouse tissue.

**Figure 5.**
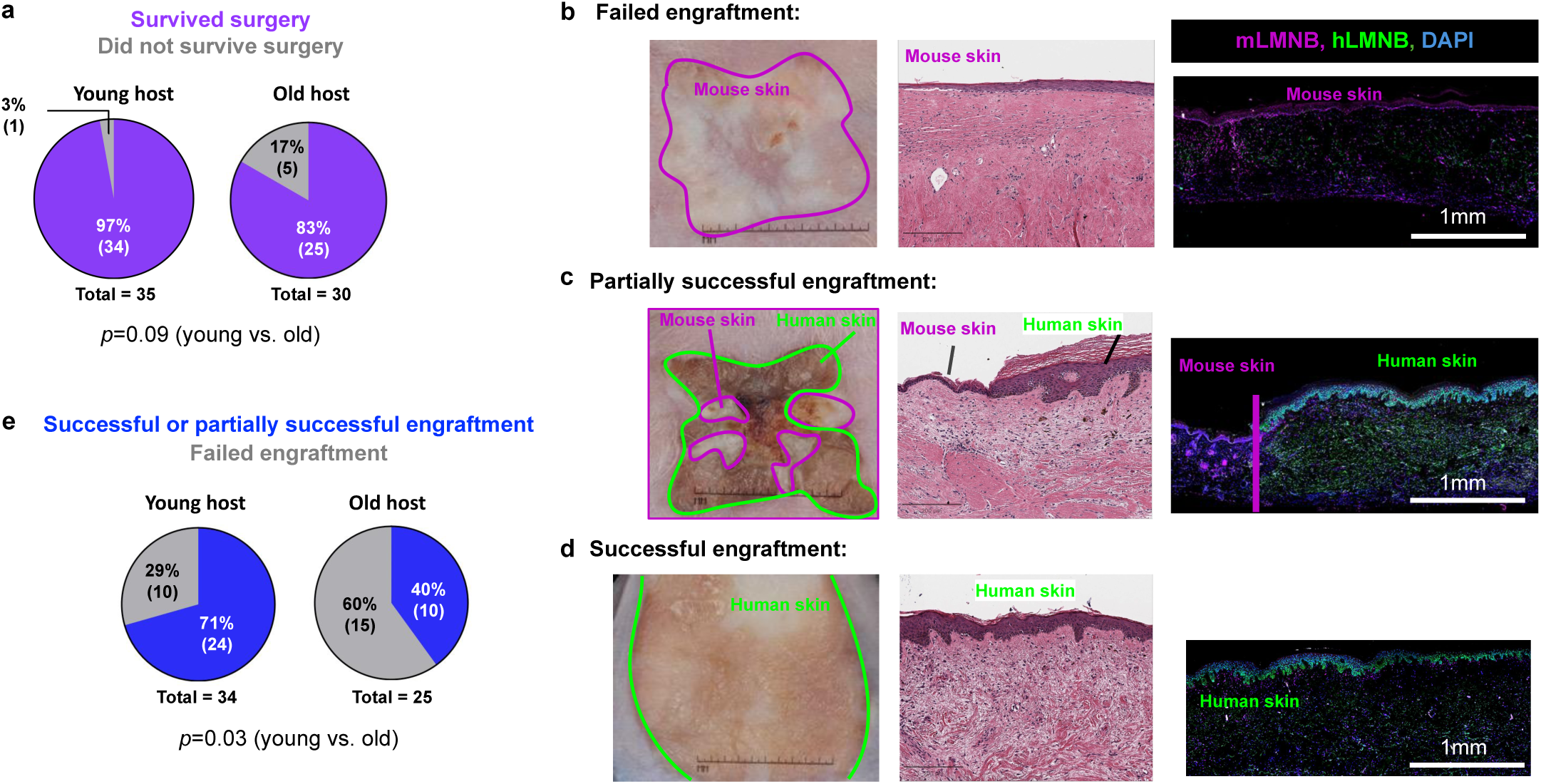
Human skin engraftment is impaired in old mice. Split-thickness human abdominal skin grafts were transplanted onto immunodeficient nude male mice (nu/nu). a. proportion of surviving animals after xenografting surgery (n = 35 young, 30 old). b. Example of a failed human skin xenograft. c. Example of partially successful human skin engraftment. d. Example of fully successful human skin engraftment. e. Proportion of successfully engrafted and failed xenografts in young and old mouse hosts (n = 34 young, 25 old). Grafts were created using skin from 7 independent donors aged 66 to 75 years, each xenografted onto multiple animals, and examined 2-3 months following surgery. Representative graft photographs, H&E stains and immunofluorescent species-specific LMNB stains are shown. P-values were calculated using Fisher’s exact test.

#### Xenografting Leads to Epidermal Restructuring

Following 2 to 3 months of graft maturation (which we had previously determined was sufficient time to discern whether the grafting was successful; data not shown), H&E sections were analyzed using the same computational epidermal pipeline described above. For each human donor, pre-graft epidermal metrics were longitudinally compared to post-graft values in both young and old mouse hosts (Figure 6a). In addition, post-graft metrics were directly compared between the two host age groups to assess whether a young systemic environment could modify or reverse age-associated structural changes in the epidermis (Figure 6b).

**Figure 6.**
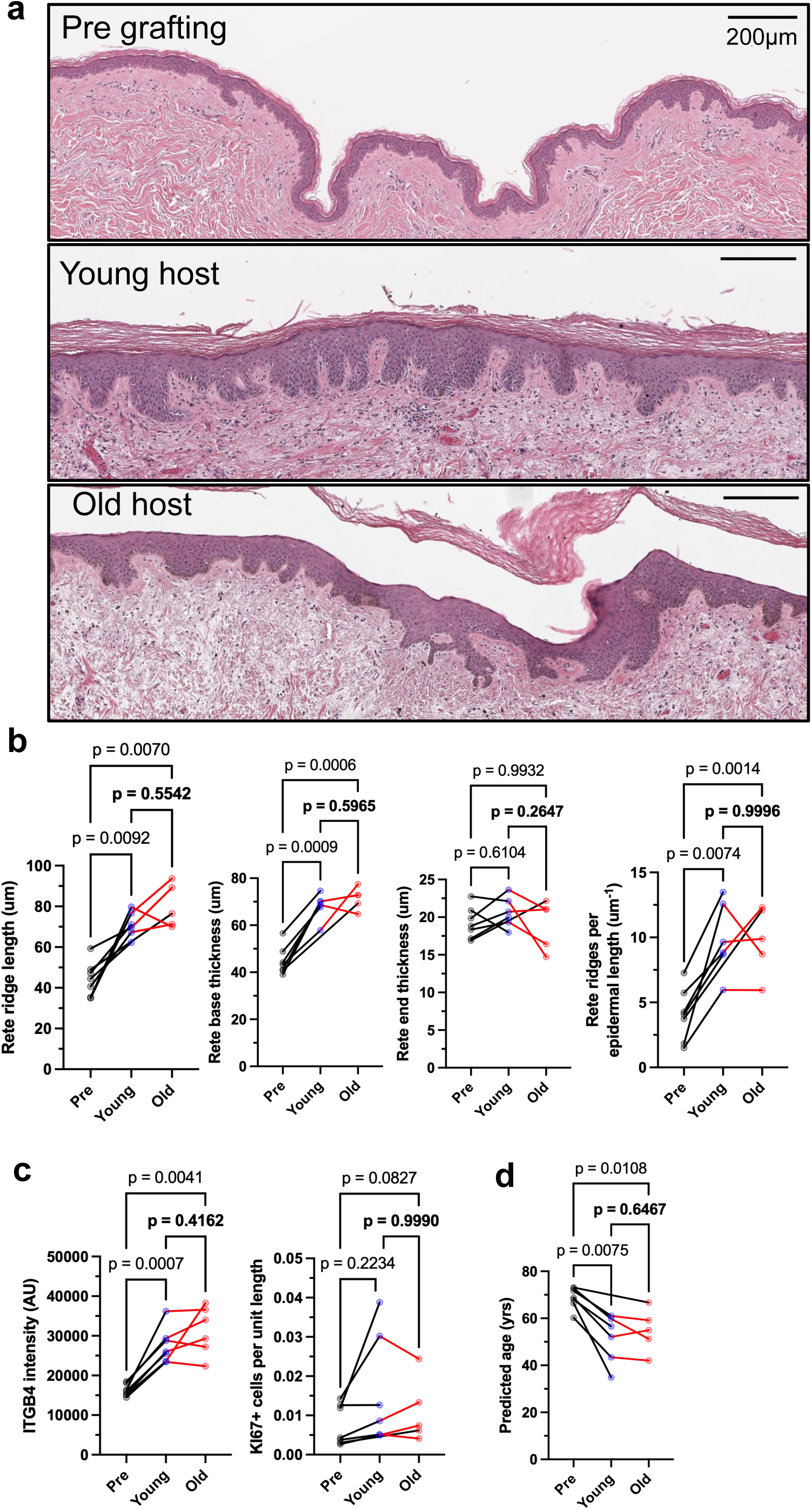
Rejuvenation of successful xenografts occurs independently of host age. a. Representative H&E sections before grafting (Pre) and after grafting into a young or an old mouse. Scale bars, 200 um. b. H&E-derived rete ridge features. c. ITGB4 intensity and Ki67+ cells per unit epidermal length. d. Model-predicted skin age. Each point is one donor (biological replicate); lines join the same donor. A donor’s graft slides in one host group were collapsed to the median (b) or mean (c, d). n (Pre, Young, Old) = 7, 6, 5 donors, except ITGB4 (7, 7, 6). P, mixed-effects analysis with the Geisser-Greenhouse correction and Tukey’s multiple comparisons test, with individual variances computed for each comparison.

Across the dataset, several features demonstrated statistically significant changes when comparing pre-graft to post-graft conditions. Notably, median rete ridge length increased significantly following grafting to both young (1.60x increase) and old (1.80x increase) mice, with no significant difference observed between the two post-graft groups. Median rete ridge base thickness also showed consistent, significant increases post-grafting, increasing by 1.52x in young hosts and 1.59x in old hosts, again with no significant difference between the young and old post-graft host groups. Median rete ridge end thickness showed no significant changes across groups. In contrast, the rete ridge dilation factor, defined as the ratio of end to base thickness, significantly decreased following grafting. The median dilation factor was reduced by 0.76x in young hosts and 0.68x in old hosts, indicating a relative thickening of the base compared to the end. Importantly, the density of rete ridges-quantified as the number of ridges per unit of epidermal length-increased markedly following grafting. Both young and old host groups exhibited similar significant increases in ridge density compared to pre-graft values (young: 2.43x; old: 2.41x), with no detectable difference between the young and old post-graft groups. Inter-rete epidermal thickness, in contrast to rete ridge metrics, showed no consistent or statistically significant changes post-grafting (data not shown).

Together, these results indicate that xenografting induces substantial epidermal restructuring that is independent of host age. Both young and old mouse environments produced similar increases in rete ridge length, base thickness, and density, accompanied by a reduction in dilation factor, reflecting a general remodeling toward a more compact and undulated epidermal architecture characteristic of younger human skin.

#### Marker Changes in the Epidermis After Xenografting

To determine whether the host environment influences molecular markers of epidermal integrity and proliferation, we quantified ITGB4 fluorescence intensity and Ki67-positive cell counts in xenografted skin derived from aged human donors (Figure 6c). As in previous analyses, ITGB4 intensity was measured within manually segmented epidermal regions, and Ki67-positive cells were manually counted and normalized by epidermal length. Comparisons were made between pre-graft samples and post-graft tissues transplanted into either young or old mice, as well as between the two post-graft groups.

For ITGB4, significant increases in expression were observed following xenografting. Mean ITGB4 intensity increased by 1.70x after grafting to young hosts and by 1.93x after grafting to old hosts, relative to pre-graft samples (Figure 6c; representative images in Supplementary Figure S9). There was no statistically significant difference between the post-graft ITGB4 levels in young versus old host environments, indicating that the restoration of ITGB4 expression was comparable in both groups. These findings suggest that xenografting alone, independent of host age, is sufficient to restore or enhance expression of basal adhesion markers in aged human epidermis. Ki67+ cells per unit epidermal length rose in every donor with matched samples (6 of 6 into young hosts and 5 of 5 into old hosts; geometric mean 1.80-fold and 2.01-fold), but the increase did not reach significance with five to six donors per group (Figure 6c, p = 0.22 and 0.083).

#### Xenografting Leads to Phenotypically Younger Skin

To assess how xenografting affects the predicted biological age of aged human skin, we applied the Random Forest regression model trained on native buttock skin to grafted samples. Predicted ages were obtained for each slice and averaged across conditions for each donor to generate estimates of age for each of the pre-graft states and post-graft states in either young or old mouse hosts. From these values, within-donor differences were calculated to evaluate the effects of xenografting on age predictions.

Across all donors with complete data, xenografting into either young or old mouse hosts resulted in a consistent decrease in predicted age (Figure 6d). Among the six donors with matched pre-graft and young-host grafts, the predicted age decreased by an average of 16.7 ± 7.7 years (mean ± SD); for the five donors with matched old-host grafts, it decreased by 14.5 ± 5.8 years. Across the four donors with grafts in both host groups, predicted age differed by 2.3 ± 4.7 years (young minus old), which was not statistically significant and was small relative to the decrease from pre-graft values. Additionally, after grafting into young or old hosts, grafts resemble never-grafted young skin across the rete ridge features that change with age (Supplementary Figure S10).

Overall, the model identified a substantial reduction in predicted biological age following xenografting, with both young and old host environments supporting this effect. Notably, the absence of a consistent or pronounced difference between young and old hosts indicates that the reduction in predicted age is driven more by the grafting process itself than by exposure to a young systemic environment. These findings are consistent with our analyses showing that key features such as ITGB4 intensity and rete ridge morphology are restored post-grafting in both host conditions.

## Discussion

The incorporation of automated, quantitative methods in this study avoids the subjective variability of manual annotation and enables high-throughput analysis across large image datasets. Our approach provides a high-resolution view of the dermal-epidermal interface by incorporating structural graph modeling of rete ridges that is well-suited to detecting subtle morphological shifts occurring during aging.

We show that aging in sun-protected skin is marked by changes at the dermal-epidermal junction consistent with weakened basal adhesion and decreased basal proliferation: ITGB4 declines with age, as do Ki67+ cells per unit epidermal length and per rete ridge. These changes align with reduced hemidesmosomal attachment and slower renewal. The data are correlative but internally consistent, indicating concerted remodeling of the epidermis with age. Interestingly, the decrease in ITGB4 expression and Ki67-positive cells was most strongly diminished in the oldest, 60+ year-old group, suggesting acceleration of epidermal aging.

The predictive modeling results demonstrate that features of sun-protected skin in this cohort contain measurable information about chronological age. Even with a relatively small dataset, the Random Forest model achieved reasonable accuracy, indicating that epidermal structure and basal layer activity encode age-related signatures detectable by computational methods. The predictive model outperformed any individual feature for predicting donor age (R² = 0.567 for the model, Figure 4b; R² = 0.38 for ITGB4, the best individual predictor, Figure 4c). Performance improvements after feature selection and polynomial expansion suggest that a limited subset of features, when allowed to interact nonlinearly, captures much of the age-related variance across individuals. The strong weighting of rete ridge end-thickness metrics highlights the morphological remodeling of the dermal-epidermal junction as a key determinant of skin aging. Supporting contributions from ITGB4 and Ki67 further indicate that basal adhesion and proliferative capacity jointly inform epidermal age. Together, these findings show that skin aging is reflected in both architectural and cellular features, and that a small, interpretable feature set is sufficient to build predictive models of epidermal age from histological data. Two recent studies also predict age from skin histology, with mean absolute errors of 7.7 years (Han et al. 2024)) and 4.9 years (Abila et al. 2026). Both cohorts are larger than ours and span different ages, and mean absolute error depends on that spread: always predicting the cohort’s median age gives an error of 14.66 years on our 41 donors but about 9.6 years on GTEx skin, so raw errors are not comparable across cohorts. When evaluating using R², which is less sensitive to data spread, the GTEx-based model reaches 0.731 compared to our 0.567. Neither study segments rete ridges or quantifies basal markers, and neither has been applied to grafted skin. Therefore, our aim was not to create a more accurate absolute age predictor, but rather to establish an interpretable epidermal readout of change within a donor.

We demonstrate that xenografting aged human skin into immunodeficient mice triggers a broad structural and molecular rejuvenation of the epidermis, reflected by both morphometric remodeling and reductions in model-predicted biological age. The observed thickening and densification of rete ridges, together with enhanced ITGB4 expression and a consistent, if not significant, rise in Ki67+ cells, imply restoration of epidermal integrity and basement membrane adhesion typical of younger skin. By the time the grafts were harvested (2-3 months after surgery), the epidermis of the grafts likely had already gone through its most active proliferative stage, which could explain why the increase in Ki67-positive cells, a marker of ongoing cell division, was modest at harvest (about twofold in every donor) and did not reach statistical significance. In contrast to prior reports (Gilhar et al. 1991; Keren et al. 2022) attributing rejuvenation primarily to systemic factors present in young but not old hosts (“young systemic factors”), we clearly show that these rejuvenating changes occurred regardless of whether the graft was hosted by a young or old mouse. Instead, our findings indicate that the principal driver of rejuvenation is the xenografting process itself, which triggers wound healing and requires revascularization and tissue integration rather than exposure to a youthful systemic milieu.

One possible explanation for these contrasting findings is the different combination of molecular features examined in the published work versus in the current study. Another important difference between the Keren et al. and our study is that in the former, histological analysis of the xenografted skin reportedly was performed 1 to 4 weeks after transplantation (Keren et al. 2022). This timepoint likely is too early to gauge whether a xenograft is successfully incorporated into the mouse, as the healing process is still ongoing and the surgical site is often covered by a scab. For this reason, we analyzed xenografted skin 2 to 3 months after surgery – a time when scabs have detached and visual examination can indicate with high reliability whether human skin is present or replaced by mouse skin tissue (Figure 5b-d; confirmed by species-specific IF staining). A failed xenograft (i.e. only scarred mouse tissue is present at the surgical site; Figure 5b) could easily be mistaken for human skin with a thin flat epidermis, resembling aged human skin, if only H&E staining is used and if gross examination of the surgical site is obscured by a scab. Previous studies using xenografted skin (Mascharak et al. 2025) may need closer scrutiny using species-specific cell markers to ensure conclusions were drawn based on human skin rather than remodeled mouse tissue. Given our finding that engraftment of human skin is significantly impaired in aged mice (Figure 5e), it is possible that such failed xenografts could mistakenly be categorized as xenografts that did not undergo rejuvenation in aged mice. Our xenograft conclusions rest on seven donors, and engraftment failed in 60% of old hosts against 29% of young hosts (Figure 5e). Two features of the design limit the effect: each donor is its own control, and the predicted-age decrease is similar in young hosts (16.7 years) and old hosts (14.5 years), so the effect cannot be from a bias acting only on the old-host arm.

By the convention of The Jackson Laboratory, mice are middle-aged from 10 to about 15 months and old from 18 to 24 months (Kevin Flurkey, Joanne M. Currer, and D.E. Harrison 2007), so our old hosts, like those of Keren et al. (14 ± 3.2 months), were grafted at the upper end of middle age. Capillary loss and reduced perfusion are documented in mouse paw skin within this window (Mesa et al. 2025), and engraftment failed more often in our old hosts (Figure 5e). Mouse trunk skin lacks rete ridges (Thompson et al. 2026), so host age can act on the graft only through the wound bed. A progeroid host would not be a suitable choice for these experiments as it could confound age with mutation-driven disease (Harkema et al. 2016).

Our results show that local graft remodeling alone (regardless of host age) can markedly reverse age-associated epidermal features, but the mechanism is unclear. One possibility is that the wound healing process induces the release of growth factors that act directly on epidermal cells (e.g. IGF, KGF). Another possibility is that the process of angiogenesis activated during wound healing rejuvenates skin by increasing skin perfusion. This interpretation is supported by recent reports of anti-aging effects of the vascular endothelial growth factor (VEGF) systemically and in the skin directly (Grunewald et al. 2021; Ichijo et al. 2022) Why graft survival is impaired in old mice is another outstanding question and could also be linked to angiogenesis. Revascularization is critical for graft incorporation (Resto and Deschler 2009) and mouse skin loses dermal capillaries with age (Ichijo et al. 2022), which would result in impaired anastomosis between human microvessels in the graft and the mouse microvessels in the wound bed. This raises the idea that modulation of the VEGF pathway could be used to increase skin graft survival in aged individuals. Interestingly, Keren et al. (Keren et al. 2022) conclude that VEGF is sufficient to induce rejuvenation of aged human skin xenografted into aged mice. However, this result is also consistent with the idea that VEGF treatment during engraftment may have led to an increased proportion of grafts with rejuvenated morphology due to better graft survival.

An important consideration is whether the observed rejuvenation phenotype is influenced by the immune characteristics of the recipient mouse strain. Although nu/nuJ mice lack functional thymus-derived T-cell immunity, they retain innate immune populations, including NK cells and macrophages, that could potentially influence the transplanted human skin through inflammatory signaling and tissue remodeling. However, similar rejuvenation effects have been reported in human skin xenografts using SCID/beige mice (Keren et al., 2022), suggesting that the phenotype is not restricted to the nu/nuJ strain. Nevertheless, because these recipient models differ in their immune deficiencies but retain aspects of innate immunity, contributions from host strain-dependent immune or other microenvironmental factors cannot be excluded. Future studies using complementary xenograft models and targeted modulation of specific immune-cell populations would help clarify whether these factors contribute to the observed rejuvenation phenotype.

Overall, the results presented here demonstrate that histological, molecular, and computational perspectives can be unified to reveal and quantify the architecture of epidermal aging and rejuvenation. Feature-based modeling provides a scalable framework for linking structure to function, enabling both discovery of age-related mechanisms and quantitative assessment of interventions. The convergence of morphometric, molecular, and predictive evidence indicates that, in this cohort, epidermal aging is a reproducible, multidimensional process that can be measured objectively and, under regenerative contexts such as xenografting, at least partially reversed.

This study is limited in scope, as it largely focuses on the epidermis. Furthermore, all donors were female and recorded by the tissue bank as Caucasian, skin phototype was not recorded, and all samples came from two sun-protected sites. Sex, ethnicity and phototype all influence skin structure and its aging (Hägg and Jylhävä 2021; Langton et al. 2019), and in an all-female cohort spanning 20 to 81 years, age is also confounded with menopausal status, which we did not record. The reference values are therefore specific to sun-protected skin of Caucasian women. We did not validate the model in an external cohort; it serves to calibrate a within-donor measurement on this cohort and is not offered as a general predictor of skin age. The study also focuses on the epidermis: dermal changes such as collagen structure, and functional readouts such as elasticity or barrier function, were not examined.

## Methods

### Human Skin Samples from Different Age Groups

Normal human skin paraffin blocks from deidentified clinical specimens were obtained from the archived tissue bank of the University of Pennsylvania SBDRC core under IRB protocol #808225. Samples were collected from sun-protected anatomical sites (buttock or abdomen) of female donors and categorized into three age groups: young (20-30 years), middle-aged (40-50 years), and old (≥60 years). Donor sex and race were recorded by the University of Pennsylvania SBDRC tissue bank and supplied with the specimens; all donors were recorded as female and Caucasian. We report race because ethnicity and phototype affect the epidermal structures measured; skin phototype was not recorded. Donors whose specimen record carried a tissue-quality note (scar, inflammation, weak staining or an unusable section; n = 9) were excluded from every analysis. The donors entering each analysis are listed in Supplementary Table S2.

### Split-thickness Human Skin Xenografting Surgery and Analysis

All animal experiments were conducted according to the Institutional Animal Care and Use Committee (IACUC)-approved protocols (#805904 assigned to GC). Discarded, unidentified human abdominal skin from seven female donors aged 66 to 75 years (Supplementary Table S5) was obtained from TenBio. Immunocompromised nu/nuJ mice were purchased from The Jackson Laboratory. Upon receipt, human skin samples were trimmed to remove subcutaneous adipose tissue and cut into 1.5 x 1.5 cm pieces. An equivalent area of mouse skin was excised to create the graft bed. Recipient mice were either young (7-10 weeks) or aged (12-15 months) at the time of grafting. Each graft was covered with sterile Vaseline gauze and secured with an elastic bandage, which was removed 10 days postoperatively. Grafts were allowed to heal for 7-11 additional weeks before evaluation and harvest. Harvested grafts were fixed in 4% paraformaldehyde (PFA) at 4°C overnight, dehydrated through graded ethanol, and embedded in paraffin.

### Immunofluorescent Staining of Tissue Sections

Paraffin-embedded tissue blocks were sectioned at 5 μm thickness. Sections were baked at 65°C for 2 hours, deparaffinized in xylene, and rehydrated through descending ethanol concentrations to water. Antigen retrieval was performed in citrate buffer (10 mM, pH 6.0) or Tris-EDTA buffer (10 mM Tris, 1 mM EDTA, pH 9.0) using a pressure cooker for 10-20 minutes. Sections were cooled to room temperature and rinsed with PBS.

After blocking with 5% normal donkey serum in PBS containing 0.1% Tween-20 for 1 hour at room temperature, sections were incubated overnight at 4°C with primary antibodies diluted in blocking buffer. Following PBS-Tween washes, sections were incubated with Alexa Fluor-conjugated secondary antibodies for 1 hour at room temperature in the dark. Nuclei were counterstained with DAPI, and slides were mounted with antifade mounting medium.

Immunofluorescence images were acquired using a Leica DM6 B microscope equipped with Leica LAS X acquisition software (Leica Microsystems). Images were acquired using HC PL APO 10×/0.45 DRY or HC PL APO 20×/0.80 DRY objectives. Fluorescence signals were detected at 460 nm (DAPI), 630 nm (RFP/CD31,TXR/mLMNB1), 525 nm (GFP/MKI67, hLMNB1), and 700 nm (Y5/ITGB4). Immunofluorescent images for ColXVII were acquired on the Akoya Vectra Polaris slide scanner at 620 nm.

### Primary Antibodies

- Anti-MKI67: eBioscience™, 48-5698-82
- Anti-ITGB4 (Integrin β4): Cell Signaling Technology, 14803S
- Anti-human LAMIN B1: SYSY Antibodies, HS-404 017
- Anti-mouse Lamin B1: SYSY Antibodies, HS-404 003
- Anti-hCD31 (PECAM-1): R&D Systems, AF806
- Anti-Collagen XVII: Abcam, ab186415

### Secondary Antibodies

- Donkey Anti-Rabbit: Thermo Fisher, Cat. No. A21207 (Alexa Fluor 594)
- Donkey Anti-Rat: Thermo Fisher, Cat. No. A21208 (Alexa Fluor 488)
- Donkey Anti-Sheep: Thermo Fisher, Cat. No. A11016 (Alexa Fluor 594)
- Rabbit anti-Rabbit: Leica, Cat. No. PV6119 + Opal tyramide signal fluorescent tag Opal 620 (Akoya, OP-001004)

### Automated Epidermal Segmentation from Whole Slide H&E Images

H&E-stained sections were scanned on an Aperio GT 450 whole-slide scanner (Leica Biosystems) at 40x magnification (0.263 µm per pixel) and supplied as pyramidal SVS files. Each image was read using the tifffile library (Gohlke). For computational efficiency, an initial downsampled version of each slide was generated by selecting the appropriate resolution level with an approximate 16x downsampling factor (4.21 µm per pixel). If no such level existed in the SVS pyramid, a zoom operation was applied via scipy to the closest larger pyramidal level to achieve the desired resolution (Virtanen et al. 2020). The downsampled image was saved for verification.

To identify tissue-containing regions, the RGB image was converted to HSV color space. Two binary masks were generated: one from the hue channel and another from the minimum intensity across channels. These masks were thresholded using Otsu’s method and combined via logical conjunction (Otsu 1979). The resulting binary mask was dilated and hole-filled to generate a preliminary tissue mask. Small connected components (<50,000 pixels) were removed. Labeled connected regions were extracted as distinct tissue slices using bounding box coordinates.

Each detected slice at downsampled resolution was mapped to its corresponding region at level 1 resolution (higher fidelity), using calculated scaling factors between the two image levels. These high-resolution slice images and corresponding binary masks were cropped and processed independently. Features were extracted at pyramid level 1, a 4x downsample of the base image, so one pixel in the feature tables is 1.052 µm; all lengths are reported in micrometres at this calibration.

For each slice, an epidermis mask was generated via color and density-based processing. First, Gaussian smoothing with a sigma of 2 was applied to each RGB channel to attenuate high frequency noise. The smoothed image was converted to both LAB and HSV color spaces. A series of thresholds were applied to remove background and non-epidermal regions, including filtering based on lightness, alpha value, hue, and saturation. The following conditional thresholds on which to mask were empirically set: L<50; L>140 (abdomen) or L>160 (grafted tissue) or L>200 (buttock); L<100 and A<150; H<110; S<40; R=G=B. The remaining tissue was deconvolved into optical density (OD) channels using a PCA-based stain separation algorithm (Ruifrok and Johnston 2001). The nuclear channel was isolated by retaining OD intensities above an empirically set threshold of 0.2, followed by Gaussian filtering and Otsu thresholding to generate a nuclear density mask. Binary morphological operations (opening, filtering small objects) were applied. The resulting binary mask is further opened (4 iterations) and uniformly filtered (10×10). Pixels with local mean > 0.5 are labelled epidermis. The lightness cut is the only segmentation parameter set per site. It follows section darkness: abdomen and graft sections were on average about 22 lightness units darker than buttock sections (Supplementary Figure S11). Supplementary Figure S11 reports the sensitivity of every age regression to this cut and to Otsu’s method applied to each section’s tissue lightness histogram, which needs no site-specific value and is provided as an option in the deposited code. Post-graft sections were analysed within a graft region outlined by hand; buttock, abdomen and pre-graft sections needed no manual annotation.

### Manual Epidermal Area Quantification and Rete Ridge Analysis

For manual epidermal area quantification, H&E-stained images were calibrated in Fiji (ImageJ, NIH, USA) according to the microscope scale bar. A 1 mm linear region of the epidermis was selected for each sample. The epidermal compartment-defined from the basal layer at the dermal-epidermal junction to the upper boundary of the stratum granulosum-was manually outlined using the polygon selection tool. The enclosed area was measured using the Analyze -> Measure function in Fiji. Rete ridges were counted along the same 1 mm linear region of the epidermis. Each downward projection of the basal layer into the dermis was defined as one rete ridge. The number of rete ridges per millimeter was recorded as the rete ridge density.

### Epidermal Graph Construction and Feature Extraction

Connected components within the epidermis mask were labeled using a connectivity of 8. These components were treated as candidate epidermal regions and referred to as “trees.” Label masks and associated region properties were extracted. A k-d tree of the tissue boundary was constructed to enable fast nearest-neighbor queries for thickness measurements (Bentley 1975).

Each epidermal tree was skeletonized using morphological thinning in scikit-image (van der Walt et al. 2014; Zhang and Suen 1984). Nodes and branches in the skeleton were defined using a 3×3 convolutional neighborhood kernel where pixels with greater than 2 neighboring pixels are considered junction nodes, and pixels with exactly 1 neighboring pixel are considered endpoints nodes. Skeleton pixels not belonging to nodes are connected component labelled as branches and associated with the two nodes they connect; only branches with exactly two nodes are retained. Nodes become vertices; branches become weighted edges (weight = branch length). The minimum spanning tree is computed via networkx (Hagberg et al. 2008). A two pass Dijkstra search is performed to obtain the longest path in the graph (Dijkstra 1959). Paths shorter than 500 pixels are discarded (slice rejected). Edges on the longest path are marked midline; all other edges are provisional rete ridges. For every provisional ridge, the endpoint node centroids were queried against the tissue boundary k-d tree to compute distances. Branches where the distal end (endpoint) was closer to the boundary than the junction (base) were discarded while remaining branches were retained as valid rete ridges.

Quantitative morphological features were computed in and aggregated in numpy for each slice and written to a cumulative CSV file via pandas (Harris et al. 2020; McKinney 2010). Extracted features included:

- Epidermal Length: Number of skeleton pixels in the midline.
- Epidermal Thickness: For every midline pixel, perpendicular distance to the nearest external tissue boundary is obtained via KD tree and doubled.
- Rete Ridge Length: Estimated as the sum of the branch length plus the distances from the base to the midline and the end to the tree boundary.
- Rete ridge base and end thickness: Distances from ridge bases and ridge ends to the external epidermal boundary (x2)
- Rete ridge Dilation Factor: Ratio of end thickness to base thickness for each ridge.
- Rete Ridge Count: Total number of validated ridges.
- Rete Ridge Density: Ridge count normalized by epidermal length.

All image processing, feature extraction, and data analysis was done in python 3.13, with the following dependencies: imagecodecs=2025.3.30, networkx=3.5, numpy=2.3.2, opencv-python=4.11.0.86, pandas=2.3.1, scikit-image=0.25.2, scipy=1.16.1, statsmodels=0.14.5, tifffile=2025.6.11

### ITGB4 and COLXVII quantification

Both ITGB4 and COLXVII are specifically expressed in basal keratinocytes. The basal epidermal layer was manually selected from each section, and mean fluorescence intensity was measured using Fiji.

### KI67 quantification

Epidermal proliferation was assessed by manual quantification of MKI67 (Ki67)-positive nuclei within the epidermis of each histological section. A defined epidermal compartment was used for all analyses. To account for regional variation in epidermal size, the total number of MKI67-positive cells per section was normalized to the measured epidermal length, yielding a proliferation index expressed as MKI67-positive cells per unit epidermal length. In addition, MKI67-positive cells located within rete ridges were specifically counted, and this value was normalized to the total number of rete ridges per section to determine the average number of proliferating cells per rete ridge.

### Random Forest Modeling for Age Prediction

To assess whether quantitative histological features could be used to predict chronological age, we trained Random Forest regression models on 41 samples with complete measurements, using 48 quantitative predictors (46 H&E morphometric features, plus basal ITGB4 intensity and Ki67+ cells per unit epidermal length). The dataset included subjects across a wide adult age range, and all samples used for modeling were from female individuals without pathological annotations. We evaluated the models with a five-fold cross validation.

Patient metadata, including age and sample identifier, was first loaded from a structured CSV file. Quantitative features from hematoxylin and eosin (H&E) stained sections, ITGB4 immunofluorescence (IF), and Ki67 IF were separately loaded and joined by a common accession ID. Each of these datasets included precomputed, slice-level aggregated features (e.g., median rete ridge thickness, mean object eccentricity, etc.) previously extracted using standardized segmentation and quantification pipelines. The final dataset consisted of numeric predictors covering a range of biological domains: epidermal morphometrics (thickness and rete ridge features), basal adhesion marker intensity (ITGB4), and proliferation marker counts (Ki67 per unit epidermal length).

We used a Random Forest regressor (sklearn.ensemble; 500 trees, random_state = 42) to predict donor age. Two models were evaluated: a full model trained on all 48 predictors, and a reduced polynomial model. For the reduced model, every step was carried out inside each training fold: a random forest was fit to the training fold, the eight predictors with the highest impurity-based importance were retained, a degree-2 polynomial expansion without a bias term was applied (8 linear, 8 squared and 28 pairwise terms; 44 predictors), and a second random forest was fit to the expanded training data. The held-out fold was used only for prediction. Performance was estimated by 5-fold cross-validation (shuffled) repeated 20 times; we report the mean absolute error (MAE) and R² as the mean over the 20 repeats with a 95% confidence interval. The full model achieved an MAE of 11.88 years (95% CI 11.61 to 12.15) and R² of 0.388 (0.358 to 0.418); the reduced polynomial model achieved 9.73 years (9.42 to 10.03) and R² of 0.567 (0.536 to 0.599) (Figure 4b). For the xenograft predictions, the reduced model was fit once to all 41 training donors with the same selection and expansion steps.

### Application of Random Forest Age Prediction Model to Xenografted Skin

To evaluate how xenografting affects predicted biological age of skin, we applied the Reduced Polynomial Random Forest Regression Model described above to the grafted human skin dataset.

Quantitative features for the xenograft experiment were extracted from H&E-stained sections, ITGB4 immunofluorescence, and Ki67 immunofluorescence. For each dataset, a unique identifier column was generated by concatenating the donor Sample number with the graft condition Age class (e.g., "Pre", "Young", or "Old"). This allowed for unambiguous merging across datasets and linkage of replicate slices from the same donor and condition. Each dataset was subset to retain only the predefined list of model-relevant features, which matched those used during training of the Random Forest model. The individual feature tables were merged using the ID column to generate a single feature matrix for inference.

To summarize prediction results at the donor level, predicted ages were averaged across slices within each (Sample, AgeClass) pair. The resulting per-sample means were pivoted into a wide-format table with one row per donor and one column for each condition (Pre, Young, and Old). From this table, within-donor differences in predicted age were calculated as:

- delta_Pre-Young: change in predicted age from pre-graft to post-graft in young mouse hosts
- delta_Pre-Old: change from pre-graft to post-graft in old hostsdelta_Young-Old: difference in predicted age between grafts placed in young versus old hosts

### Statistical Analysis

Associations between each feature and donor age were tested by ordinary least-squares regression with one value per donor (the median over a donor’s H&E sections; the mean over CD31 regions of interest and sections). For every feature we report the slope with its 95% confidence interval, Pearson r as the effect size, R², the two-sided p-value and a Benjamini-Hochberg q-value computed within each family of tests, a family being one modality at one site (Supplementary Table S3, Supplementary Table S4). Features with q < 0.05 are described as significant, and associations with R² < 0.10 are treated as weak whatever their p-value. The biological replicate is the donor: technical replicates (sections, regions of interest, grafts) were collapsed to one value per donor before testing, and n in the legends counts donors. Pre-graft versus post-graft comparisons (Figure 6) used a mixed-effects analysis with the Geisser-Greenhouse correction and Tukey’s multiple comparisons test in Prism, and survival and engraftment proportions (Figure 5) used Fisher’s exact test.

## Supporting information

Supplementary_Table_HE_features

Supplementary_Table_IF_features

## Conflict of Interest Statement

The authors declare no competing interests.

## Data Availability Statement

The image-processing pipeline, per-section feature tables and analysis scripts are available at https://github.com/calico/skin-aging-xenograft. Whole-slide and immunofluorescence images with de-identified metadata are available on BioImage Archive at https://doi.org/10.6019/S-BIAD4417.

## Acknowledgments

Support for this work was provided by the Penn Skin Biology and Diseases Resource-based Center, funded by NIH/NIAMS grant P30-AR069589 (PI: Grice), the Department of Dermatology of University of Pennsylvania Perelman School of Medicine, The Edwin & Fannie Gray Hall Center for Human Appearance at University of Pennsylvania Medical Center and through a sponsored research agreement from Calico Life Sciences.

## Supplementary Materials

### Supplementary Methods

#### CD31 quantification

CD31 immunofluorescence images were analyzed to quantify microvasculature within the papillary dermis. For each histological section, the papillary dermis was manually segmented as a 100 micron deep region of the dermis directly below the dermal-epidermal junction to define the region of interest. Within this region, CD31-positive signal was identified using intensity-based thresholding and converted to a binary representation. Connected components analysis was used to segment individual CD31-positive structures. Quantitative features were extracted from segmented objects and aggregated at the section level. In parallel, microvasculature density was calculated by measuring the total CD31-positive area and expressing it as a percentage of the annotated papillary dermal area. For each sample, five papillary dermis regions were analyzed, and values were averaged to obtain a mean microvascular density per sample for comparison across age groups.

#### CD31 microvascular morphometrics in the papillary dermis

CD31-positive objects were segmented within five manually annotated papillary dermis regions per section of buttock skin from the age-stratified cohort (n = 30 donors, 31 sections; Supplementary Figure S3a). We measured the CD31-positive area fraction, the object density, and distributional statistics of object area, perimeter, solidity, and major and minor axis length, 43 features in all (Supplementary Table S4). Each feature was regressed on donor age, and p-values were corrected within the 43-feature family by the Benjamini-Hochberg procedure. Eleven features had uncorrected p < 0.05, among them mean solidity (R² = 0.14, p = 0.045) and mean minor axis length (R² = 0.14, p = 0.044), but none remained significant after correction (smallest q = 0.087). CD31 object density and area fraction did not change with age (p = 0.33 and 0.79). The sections were exported in two batches at different pixel sizes (2.049 and 1.150 µm per pixel) and with different intensity thresholds, so dimensioned features are not comparable between batches; no feature reaches q < 0.05 within either batch or with batch as a covariate. We therefore treat these data as an exploratory screen that does not establish an age-related change in papillary microvascular geometry. CD31 features were not used in the age model.

#### Solidity and the epidermal markers

Within the CD31 cohort (Supplementary Figure S3), mean solidity correlated with basal ITGB4 intensity (r = −0.38, p = 0.040, n = 29) but not significantly with Ki67+ cells per unit epidermal length (r = −0.34, p = 0.09, n = 26, Supplemental Figure S4). Because donor age could drive all three measurements, we also computed partial correlations controlling for age. Neither was significant (ITGB4: r = −0.25, 95% CI −0.57 to 0.13, p = 0.19; Ki67: r = −0.17, 95% CI −0.53 to 0.24, p = 0.41; q = 0.39 and 0.41 across the two tests). The result was the same for Ki67+ cells per rete ridge, with staining batch as an additional covariate, and with the nine donors carrying a tissue-quality note included (n = 36 and 34). Among donors of the same age, those with more solid vessels did not have detectably lower ITGB4 or Ki67, although 29 and 26 donors cannot rule out a modest association.

### Supplementary Tables

**Supplementary Table S1.**
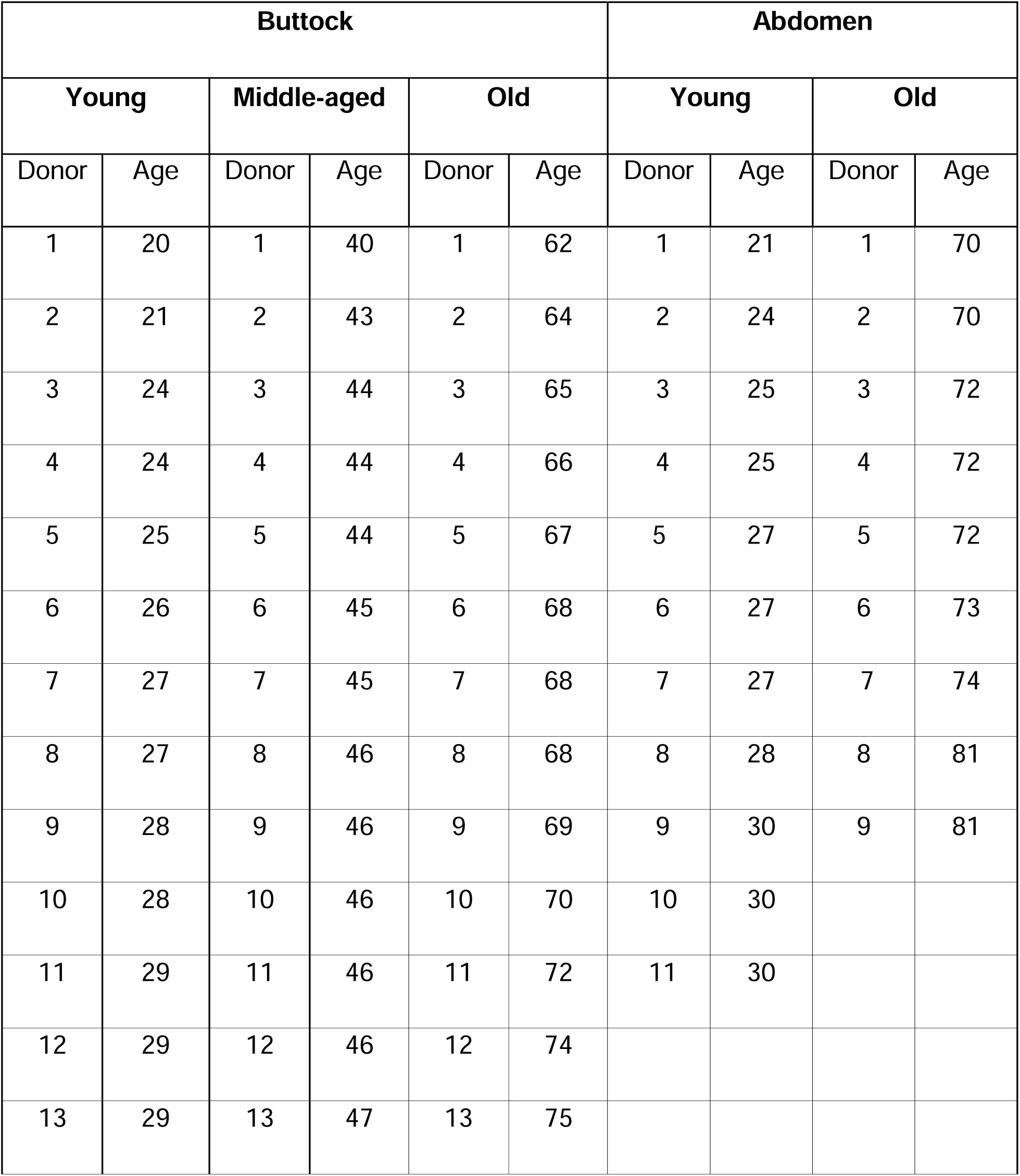
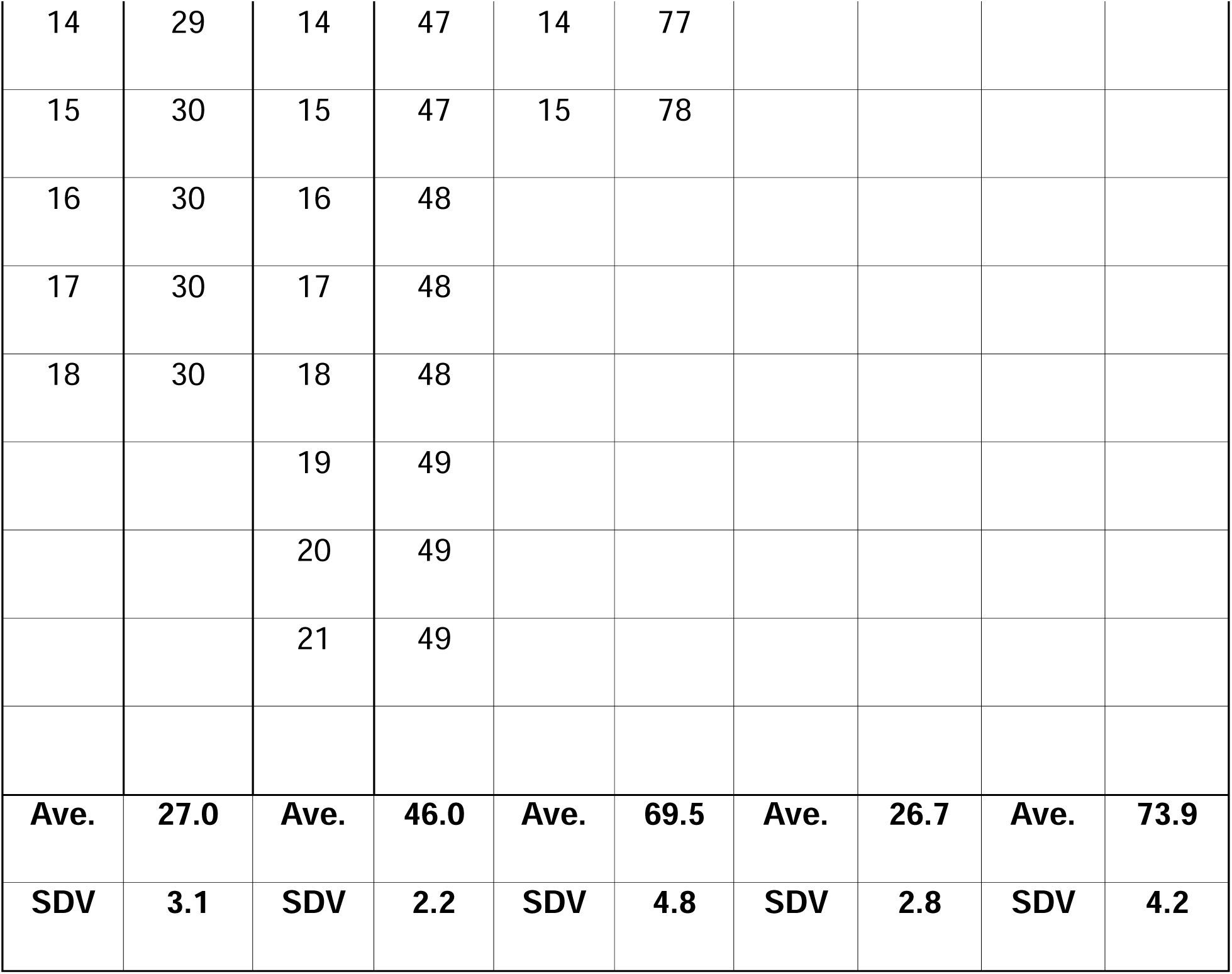
Cross-sectional skin characterization sample list.

| **Buttock** |  |  |  |  |  | **Abdomen** |  |  |  |
|---|---|---|---|---|---|---|---|---|---|
| **Young** |  | **Middle-aged** |  | **Old** |  | **Young** |  | **Old** |  |
| Donor | Age | Donor | Age | Donor | Age | Donor | Age | Donor | Age |
|---|---|---|---|---|---|---|---|---|---|
| 1 | 20 | 1 | 40 | 1 | 62 | 1 | 21 | 1 | 70 |
| 2 | 21 | 2 | 43 | 2 | 64 | 2 | 24 | 2 | 70 |
| 3 | 24 | 3 | 44 | 3 | 65 | 3 | 25 | 3 | 72 |
| 4 | 24 | 4 | 44 | 4 | 66 | 4 | 25 | 4 | 72 |
| 5 | 25 | 5 | 44 | 5 | 67 | 5 | 27 | 5 | 72 |
| 6 | 26 | 6 | 45 | 6 | 68 | 6 | 27 | 6 | 73 |
| 7 | 27 | 7 | 45 | 7 | 68 | 7 | 27 | 7 | 74 |
| 8 | 27 | 8 | 46 | 8 | 68 | 8 | 28 | 8 | 81 |
| 9 | 28 | 9 | 46 | 9 | 69 | 9 | 30 | 9 | 81 |
| 10 | 28 | 10 | 46 | 10 | 70 | 10 | 30 |  |  |
| 11 | 29 | 11 | 46 | 11 | 72 | 11 | 30 |  |  |
| 12 | 29 | 12 | 46 | 12 | 74 |  |  |  |  |
| 13 | 29 | 13 | 47 | 13 | 75 |  |  |  |  |
| 14 | 29 | 14 | 47 | 14 | 77 |  |  |  |  |
| 15 | 30 | 15 | 47 | 15 | 78 |  |  |  |  |
| 16 | 30 | 16 | 48 |  |  |  |  |  |  |
| 17 | 30 | 17 | 48 |  |  |  |  |  |  |
| 18 | 30 | 18 | 48 |  |  |  |  |  |  |
|  |  | 19 | 49 |  |  |  |  |  |  |
|  |  | 20 | 49 |  |  |  |  |  |  |
|  |  | 21 | 49 |  |  |  |  |  |  |
| **Ave.** | **27.0** | **Ave.** | **46.0** | **Ave.** | **69.5** | **Ave.** | **26.7** | **Ave.** | **73.9** |
| **SDV** | **3.1** | **SDV** | **2.2** | **SDV** | **4.8** | **SDV** | **2.8** | **SDV** | **4.2** |

**Supplementary Table S2.** Donors in each analysis.

| Analysis | Site | Donors analysed | Age years (mean +/- SD) | Technical replicates |
|---|---|---|---|---|
| H&E morphometrics | buttock | 54 | 20 to 78 (46.2 ± 17.0) | 2 to 8 tissue sections per donor (median 4); median per donor |
| H&E morphometrics | abdomen | 20 | 21 to 81 (48.0 ± 24.3) | 1 to 8 tissue sections per donor (median 2); median per donor |
| ITGB4 immunofluorescence | buttock | 52 | 20 to 78 (45.7 ± 17.0) | 1 section per donor |
| Ki67 immunofluorescence | buttock | 43 | 20 to 78 (47.5 ± 17.8) | 1 section per donor |
| CD31 immunofluorescence (exploratory) | buttock | 30 | 20 to 78 (52.6 ± 17.9) | 1 to 2 stained sections per donor, five regions of interest each; regions pooled per section, sections averaged per donor |
| Age model, training set | buttock | 41 | 20 to 78 (46.9 ± 17.8) | 1 row per donor after the per-modality collapse |
| Xenograft donors, pre-graft skin | abdomen | 7 | 66 to 75 (70.0 ± 2.8) | 1 slide per donor |
| Xenograft donors, young-host grafts | abdomen | 6 |  | 2 to 6 grafts per donor (H&E); Ki67 1 to 5; ITGB4 1 to 2; mean per donor and host group; ITGB4 measured in 7 donors |
| Xenograft donors, old-host grafts | abdomen | 5 |  | 1 to 3 grafts per donor (H&E); Ki67 1 to 3; ITGB4 1; mean per donor and host group; ITGB4 measured in 6 donors |

**Supplementary Table S3. All H&E-derived features**

Provided as an additional file, Supplementary_Table_HE_features.csv. Linear regression of each of 47 H&E features on donor age in buttock (n = 54 donors) and abdominal (n = 20 donors) skin, one value per donor (the median over its sections). Columns give the feature’s name in the manuscript and in the deposited feature tables; its unit (lengths in micrometres at 1.052 µm per pixel); the intercept (fitted value at age 0); the slope per year with its 95% confidence interval; Pearson r with its 95% confidence interval (Fisher z) and R²; the two-sided p for zero slope; and the Benjamini-Hochberg q within each site’s 47 tests. Features with q < 0.05 are significant after correction; we treat R² < 0.10 as weak whatever its p-value.

**Supplementary Table S4. All immunofluorescence-derived features**

Provided as an additional file, Supplementary_Table_IF_features.csv. Linear regression on donor age of ITGB4 intensity (n = 52 donors), Ki67-positive cells per unit epidermal length and per rete ridge (n = 43) and 43 CD31 features (n = 30) in buttock skin, one value per donor; columns as in Supplementary Table S3, with q within each marker. The main columns exclude the nine donors with a tissue-quality note; the columns marked "all donors" repeat each test with them included (ITGB4 n = 59, Ki67 n = 51, CD31 n = 39). ITGB4 intensity is in arbitrary units and CD31 lengths and areas in image pixels. Five CD31 50th-percentile columns duplicate the medians and are listed because both are in the deposited feature set.

**Supplementary Table S5.**
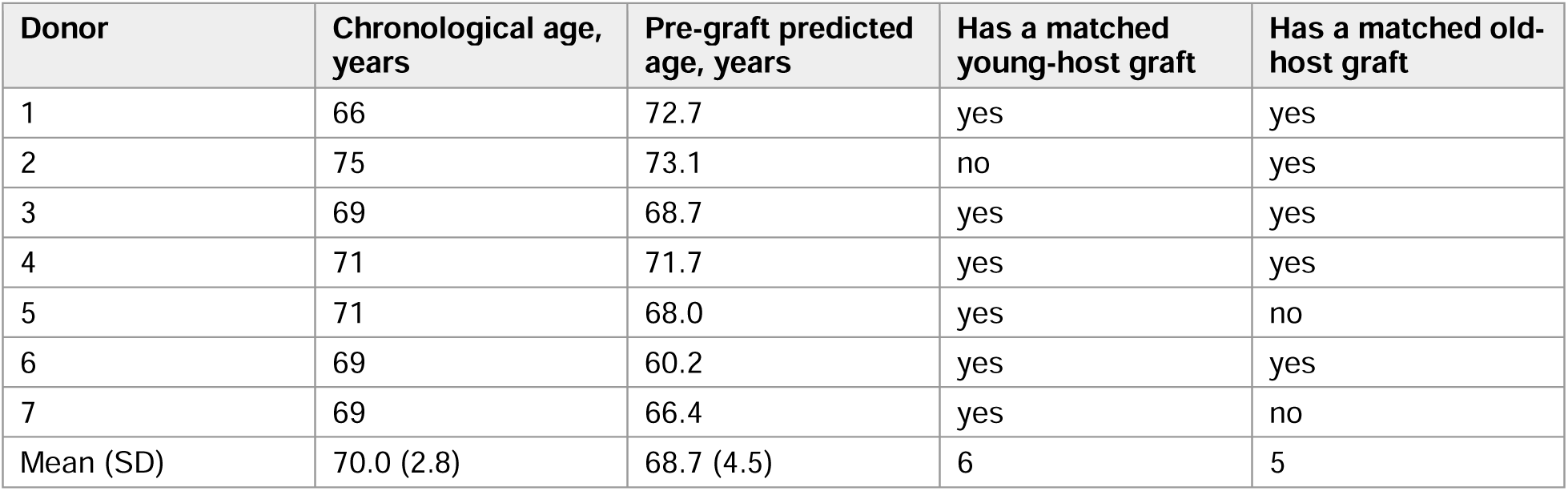
Chronological and predicted ages of xenograft donors.

| **Donor** | **Chronological age, years** | **Pre-graft predicted age, years** | **Has a matched young-host graft** | **Has a matched old-host graft** |
|---|---|---|---|---|
| 1 | 66 | 72.7 | yes | yes |
| 2 | 75 | 73.1 | no | yes |
| 3 | 69 | 68.7 | yes | yes |
| 4 | 71 | 71.7 | yes | yes |
| 5 | 71 | 68.0 | yes | no |
| 6 | 69 | 60.2 | yes | yes |
| 7 | 69 | 66.4 | yes | no |
| Mean (SD) | 70.0 (2.8) | 68.7 (4.5) | 6 | 5 |

---

**Supplementary Figure S1.**
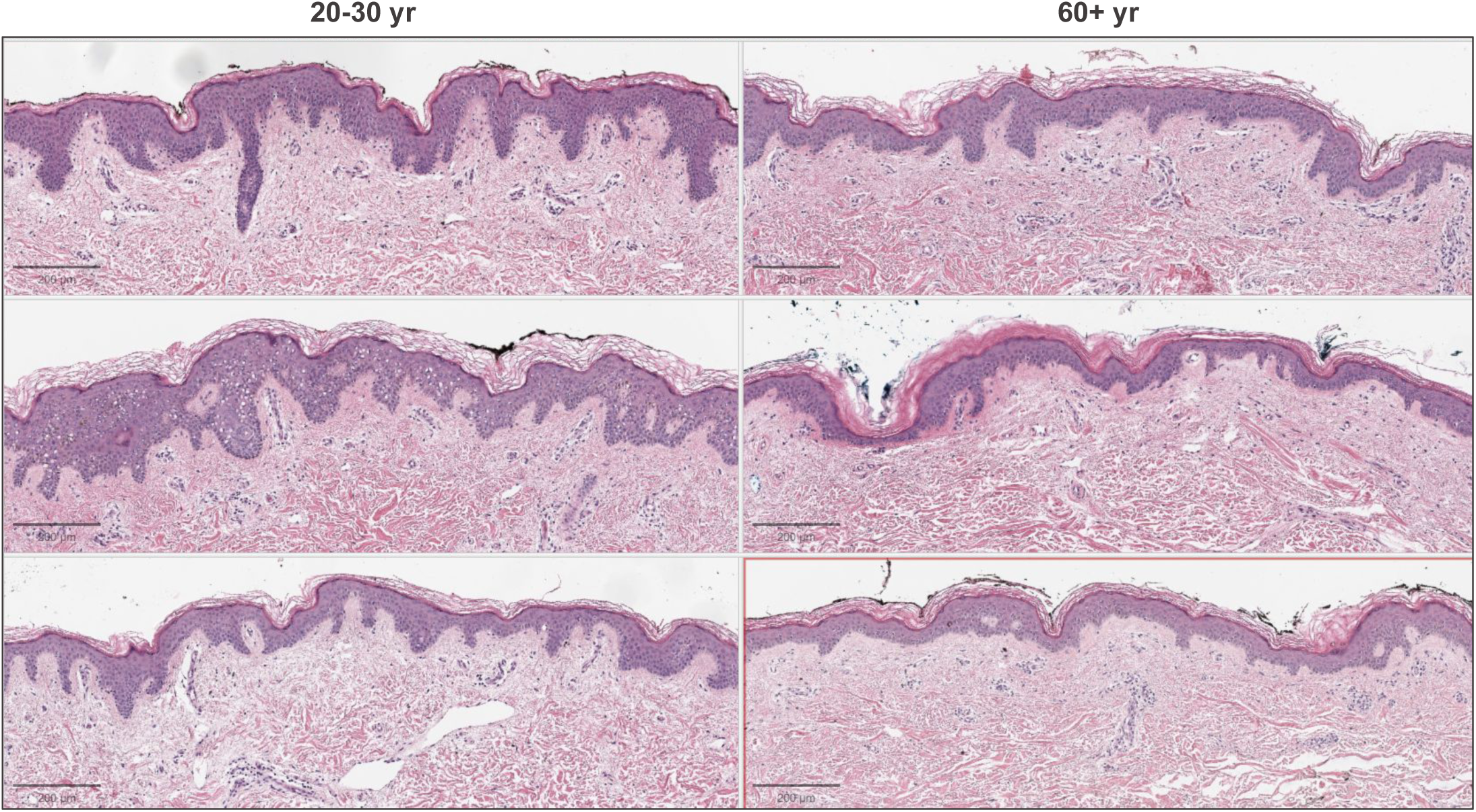
Representative images of young and old H&E-stained buttock skin. Representative H&E sections of buttock skin from three young (20-30 years, left) and three old (60 years and over, right) donors. Scale bars, 200 um.

**Supplementary Figure S2.**
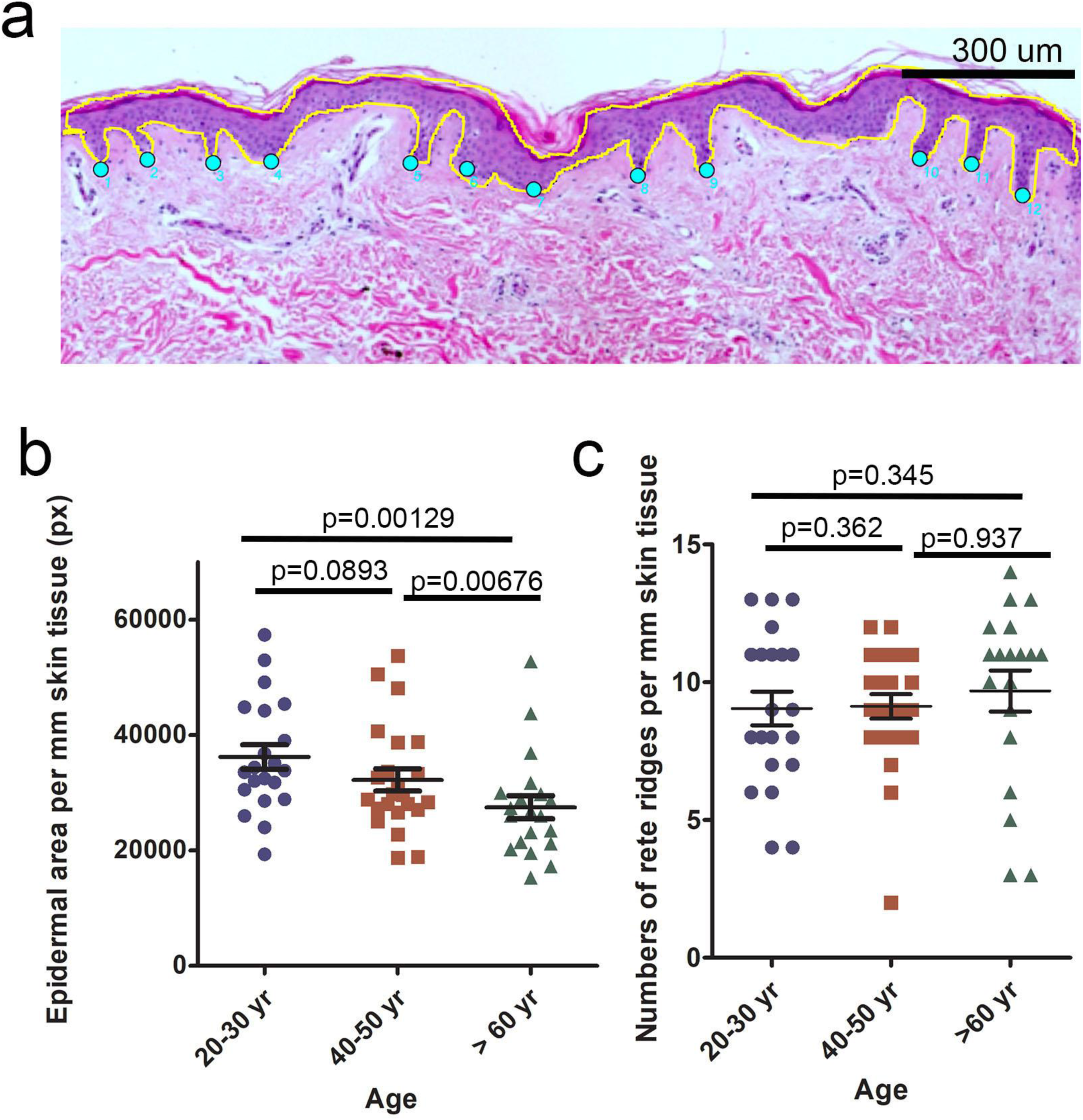
Manual quantification of epidermal thickness and rete ridge density. a. Representative H&E-stained sections of paraffin-embedded human skin tissue from young, middle-aged and old grops. The epidermal compartment was digitally delineated and measured, and rete ridges were identified as downward projections of the epidermis into the dermis. b. Quantification of epidermal area across age groups. The delineated epidermal region was measured using Fiji analysis software. c. Quantification of rete ridge number across age groups. Rete ridges were annotated in each section and counted across predefined fields of view. Counts were averaged per sample to generate group means. Data are presented as mean ± SEM (n = 19-21 per group, 1 field analyzed per sample). Error bars represent SEM. Statistical comparisons among groups were performed using one-way ANOVA test. p < 0.05 was considered statistically significant.

**Supplementary Figure S3.**
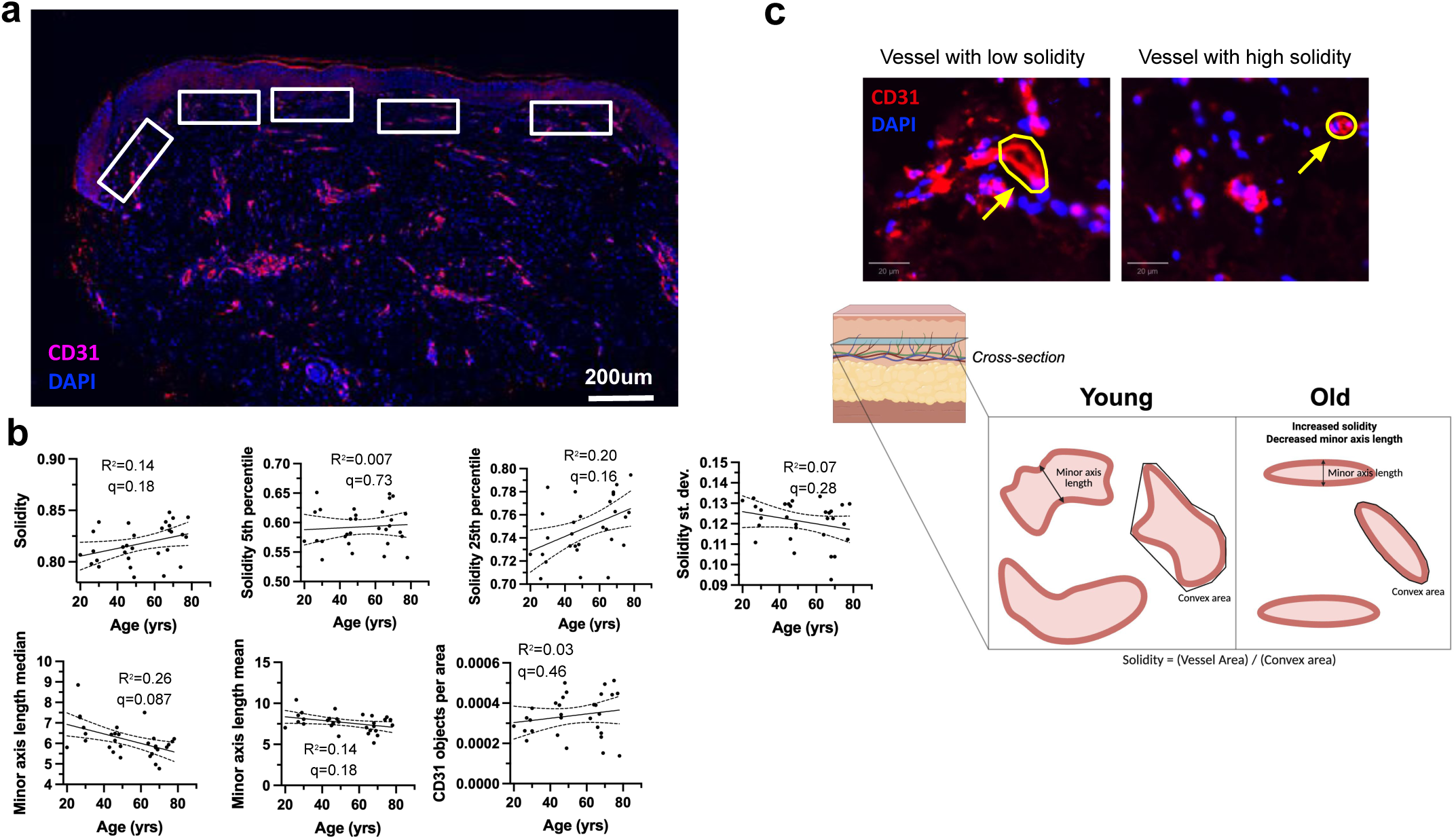
Exploratory analysis of papillary capillary morphology and age. a. Representative images of CD31 staining for endothelial cells in the papillary dermis. For quantification, five rectangular regions of interest were selected across the papillary dermis in each stained section. b. Selected CD31 features against donor age (n = 30 donors; five regions of interest per section, sections averaged per donor). Lines are least-squares fits with 95% confidence bands; R², p and the Benjamini-Hochberg q across the 43 CD31 features are shown. No feature has q < 0.05. c. Model of young and old skin papillary capillary morphometrics.

**Supplementary Figure S4.**
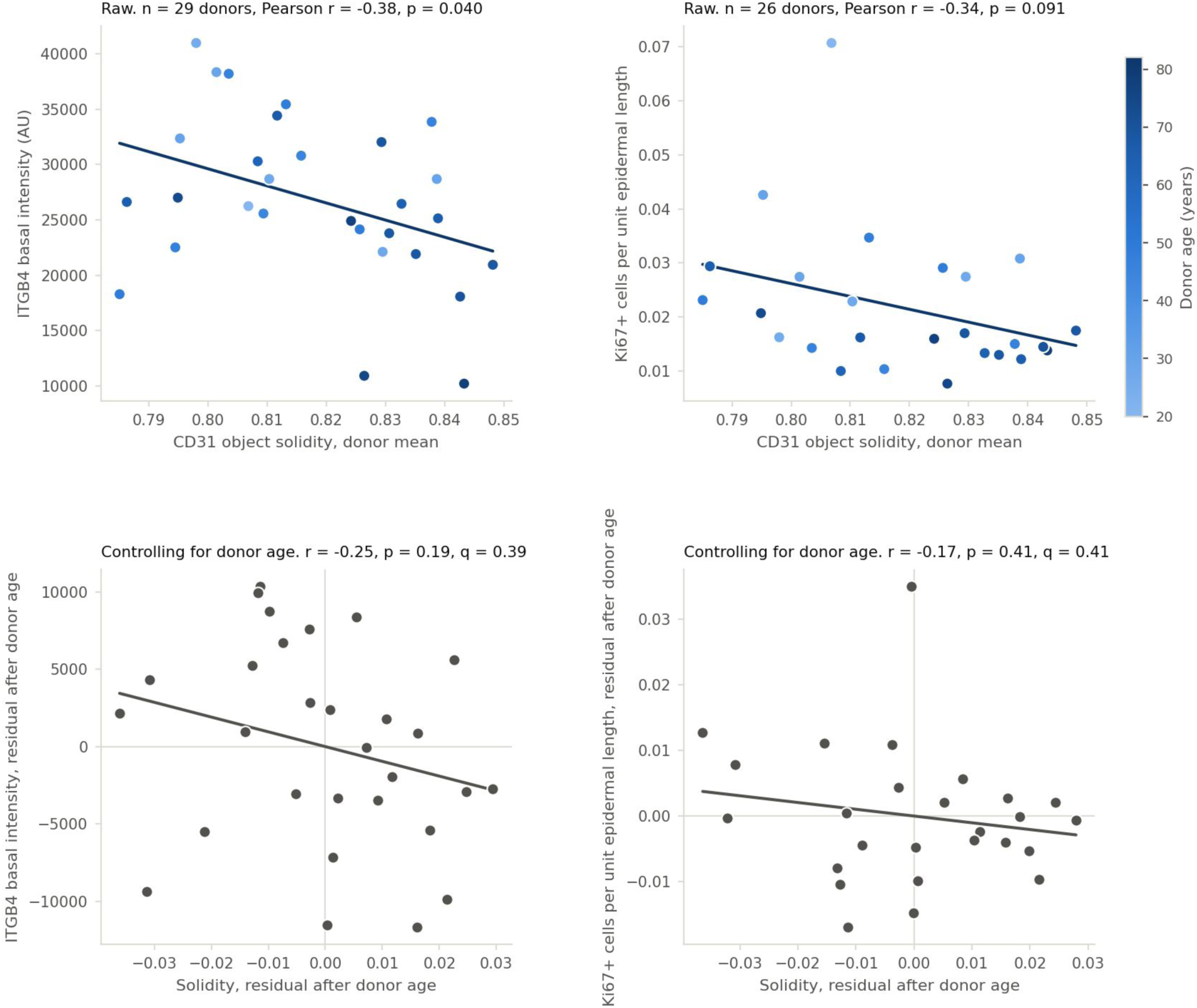
CD31 vessel solidity and the epidermal ageing markers. Each point is one donor. (a) Mean CD31 object solidity against basal ITGB4 intensity (n = 29; Pearson r = −0.38, p = 0.040). (b) Mean solidity against Ki67+ cells per unit epidermal length (n = 26; r = −0.34, p = 0.091). In (a, b), color shows donor age and lines are least-squares fits. (c, d) The same pairs after regressing each variable on donor age and plotting the residuals (partial correlations): r = −0.25, p = 0.19, q = 0.39 for ITGB4 and r = −0.17, p = 0.41, q = 0.41 for Ki67. q values are Benjamini-Hochberg adjusted across the two tests.

**Supplementary Figure S5.**
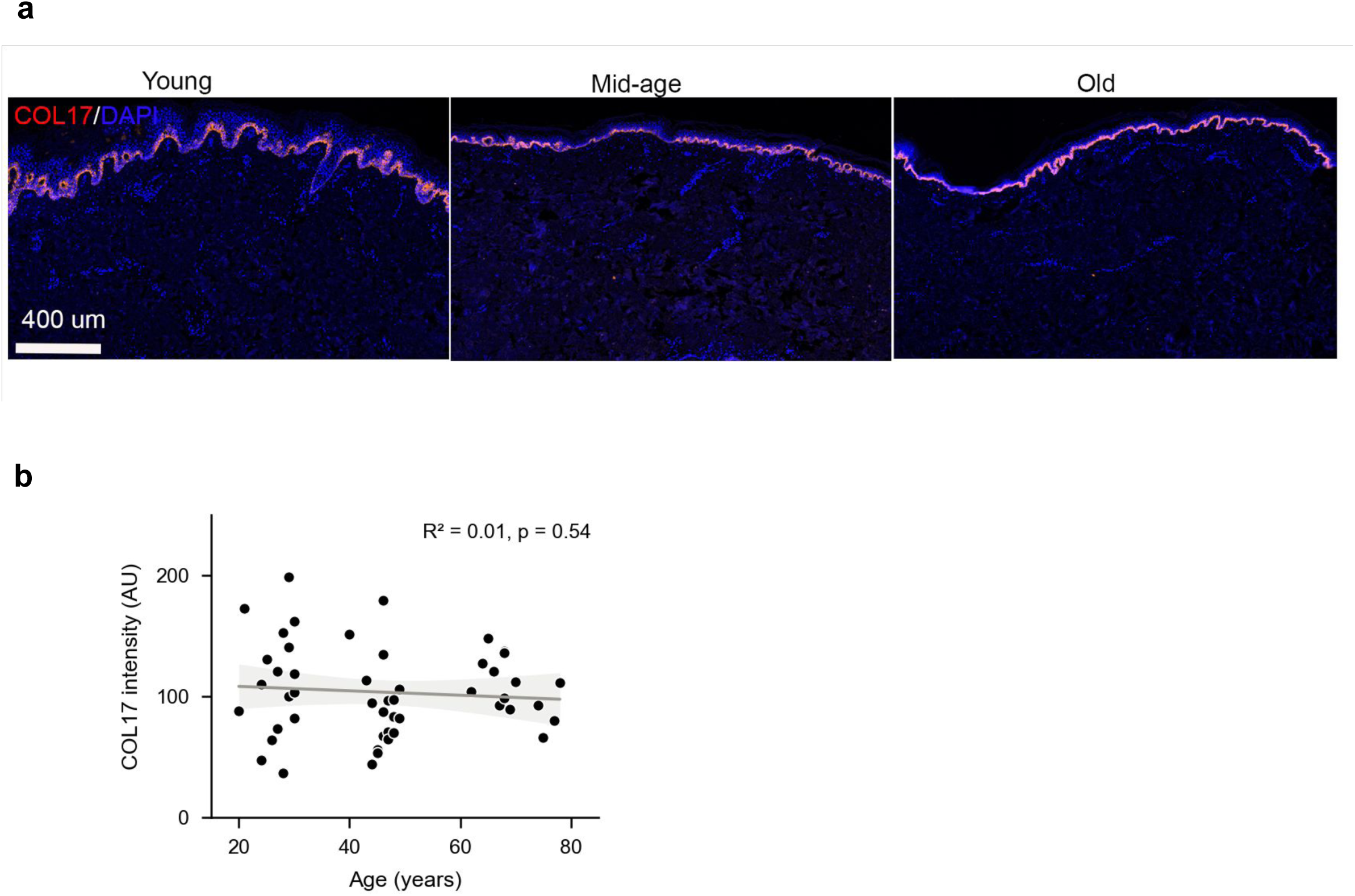
Collagen 17a levels do not correlate with age. a.Representative immunofluorescence images of skin sections from different age groups showing COL17 (red) expression predominantly in the basal layer of the epidermis. Nuclei are counterstained with DAPI (blue). Scale bar, 400 μm. b. Each point is one donor (n = 52, aged 20 to 78 years). Simple linear regression with R² and p shown; the line is the fit with its 95% confidence interval. AU, arbitrary units.

**Supplementary Figure S6.**
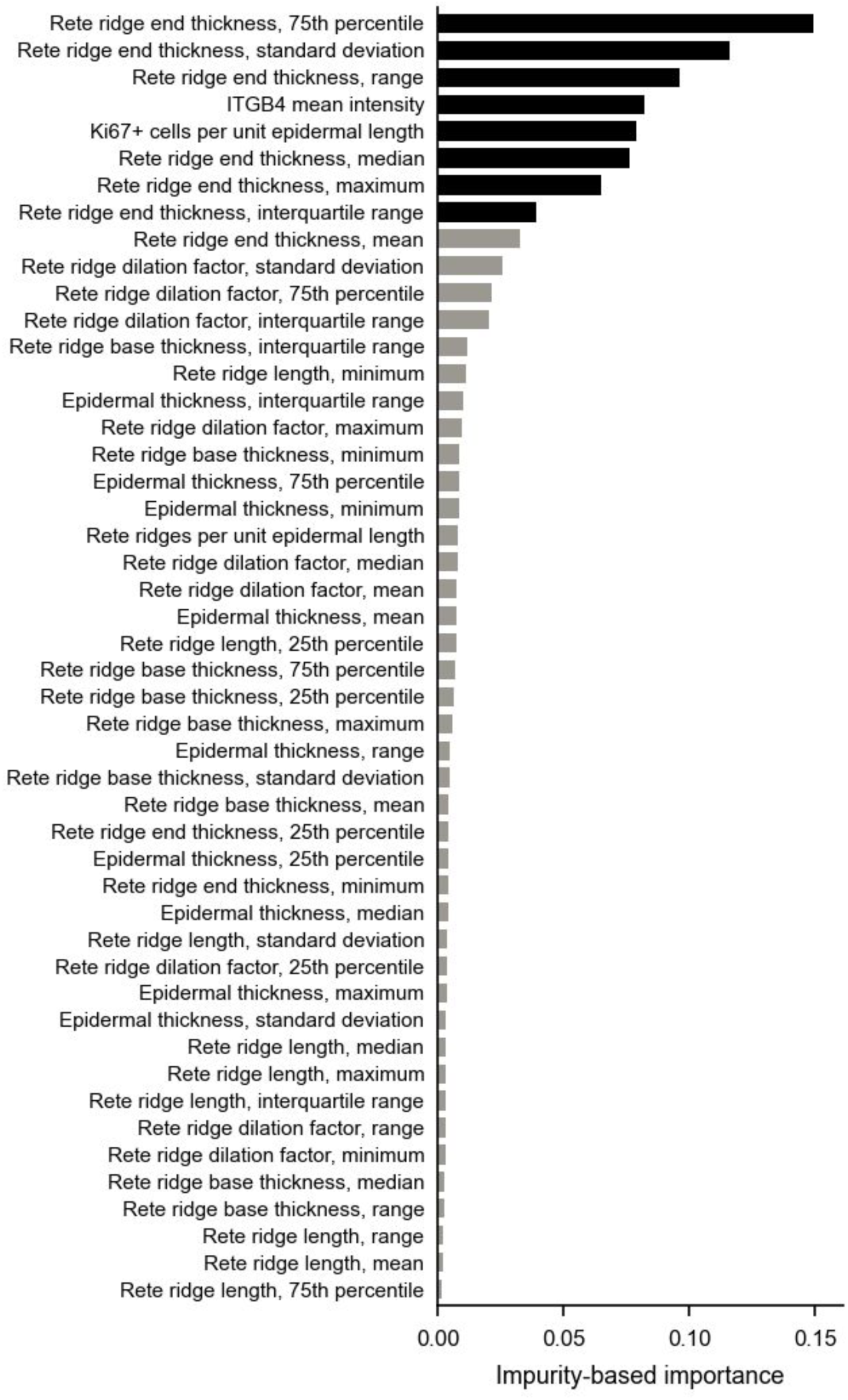
Feature importance in the full-feature random forest. Impurity-based importance of the 48 candidate features in a random forest fit to all 41 training donors. This ranking selected the eight features (black) of the final reduced model used for the xenograft predictions. The cross-validated performance in Figure 4b was estimated with this selection repeated inside each training fold. n = 41 donors, as in Figure 4b.

**Supplementary Figure S7.**
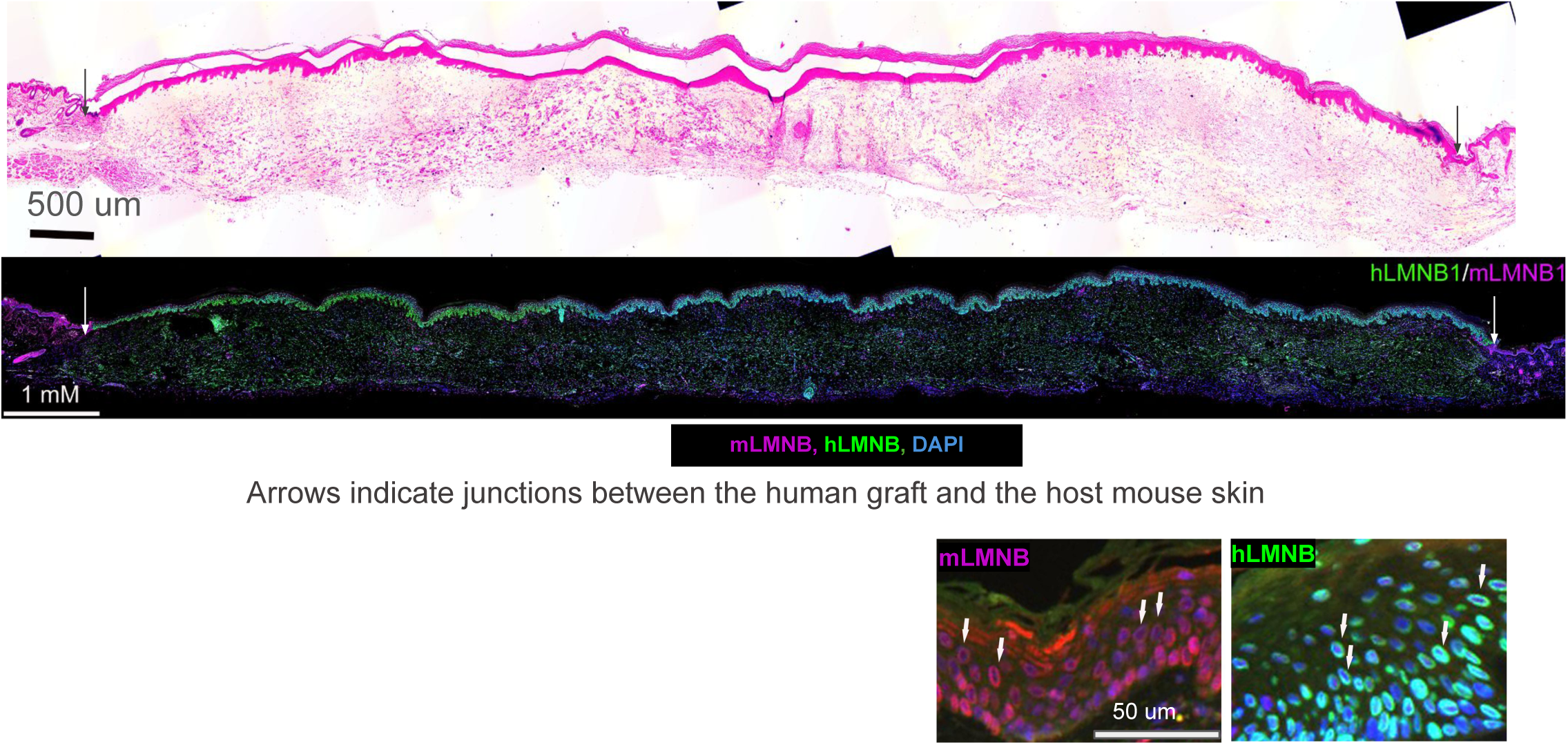
Example of a successful human skin xenograft. Representative images of the 34 successful or partially successful engraftment of human skin as confirmed by H&E staining. Immunofluorescence staining with mouse- and human-specific LAMIN antibodies was performed to distinguish mouse and human cells within the grafted skin. Arrows indicate junctions between the human graft and the host mouse skin.

**Supplementary Figure S8.**
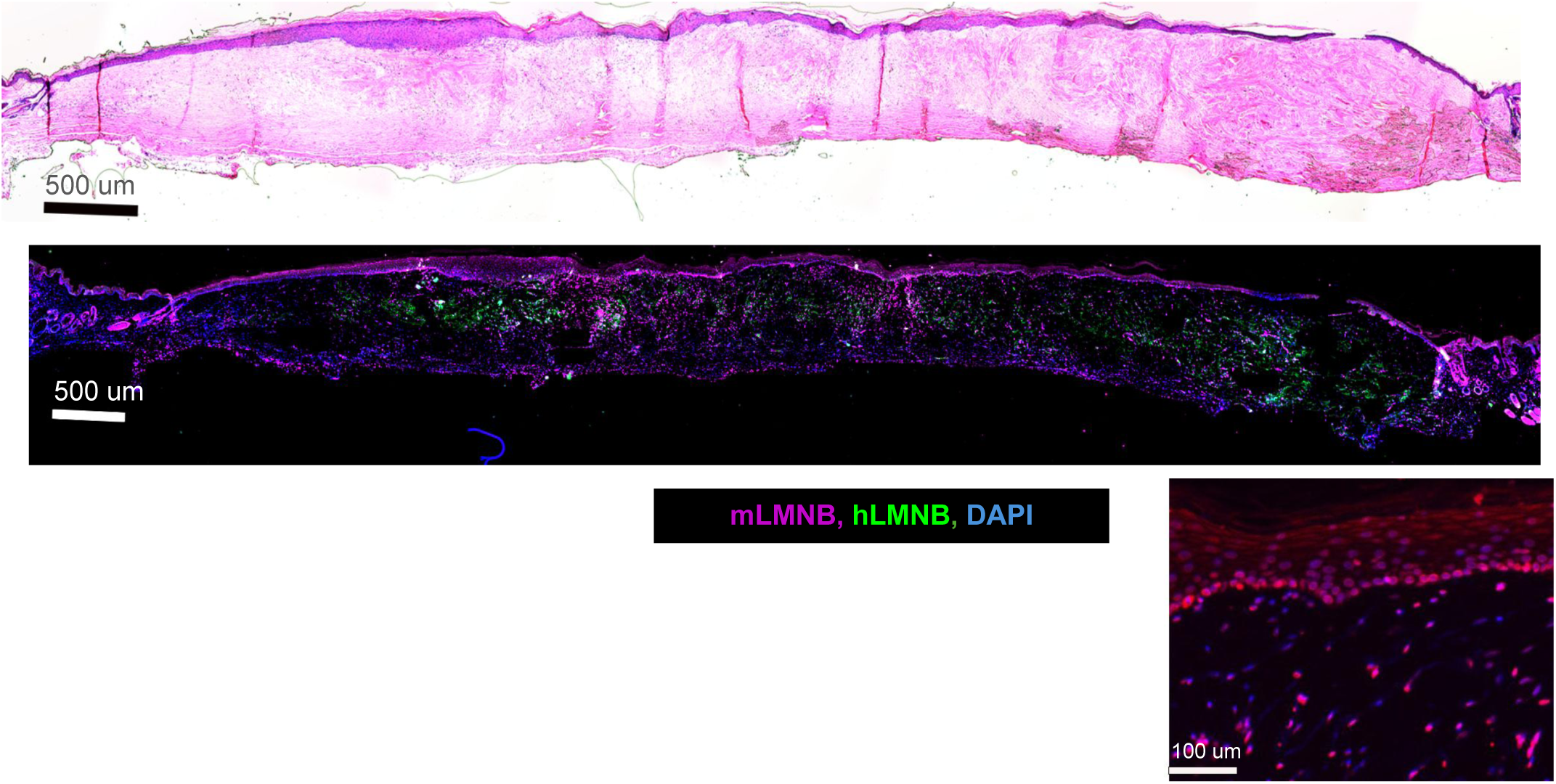
Example of a failed human skin xenograft. Representative images of the 25 failed human skin grafts. H&E staining demonstrates the absence of organized human epidermal structures, indicating graft failure. Immunofluorescence staining with mouse- and human-specific LAMIN antibodies was used to verify the cellular origin within the graft area.

**Supplementary Figure S9.**
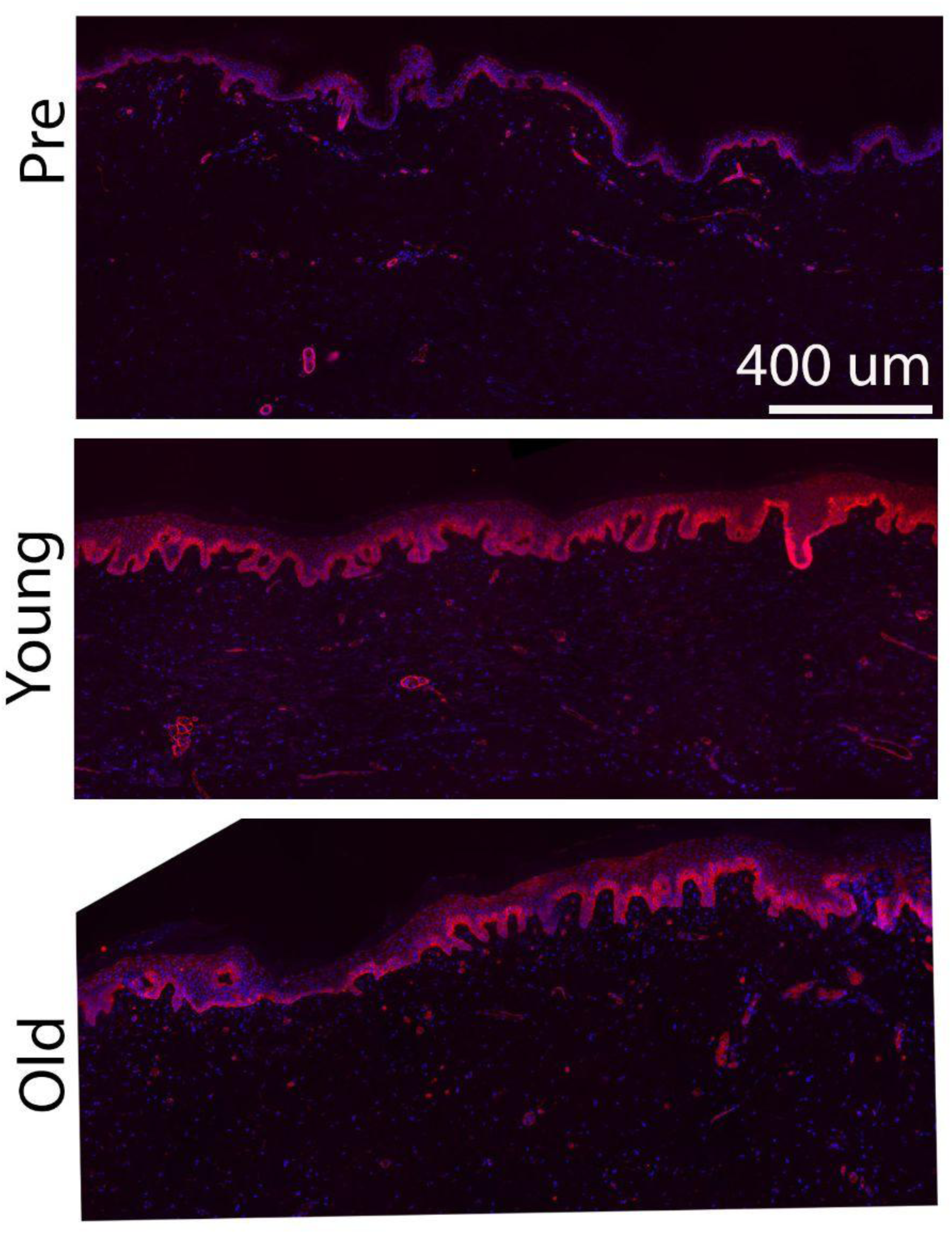
Immunofluorescence staining of Integrin β4 (ITGB4) in tissue grafts. Representative images of ITGB4 staining (red) across three experimental conditions: pre-graft baseline control (n=7), graft transplanted into a young mouse host (n=7), and graft transplanted into an old mouse host (n=6). Nuclei are counterstained with DAPI (blue). Scale bar = 400 um.

**Supplementary Figure S10.**
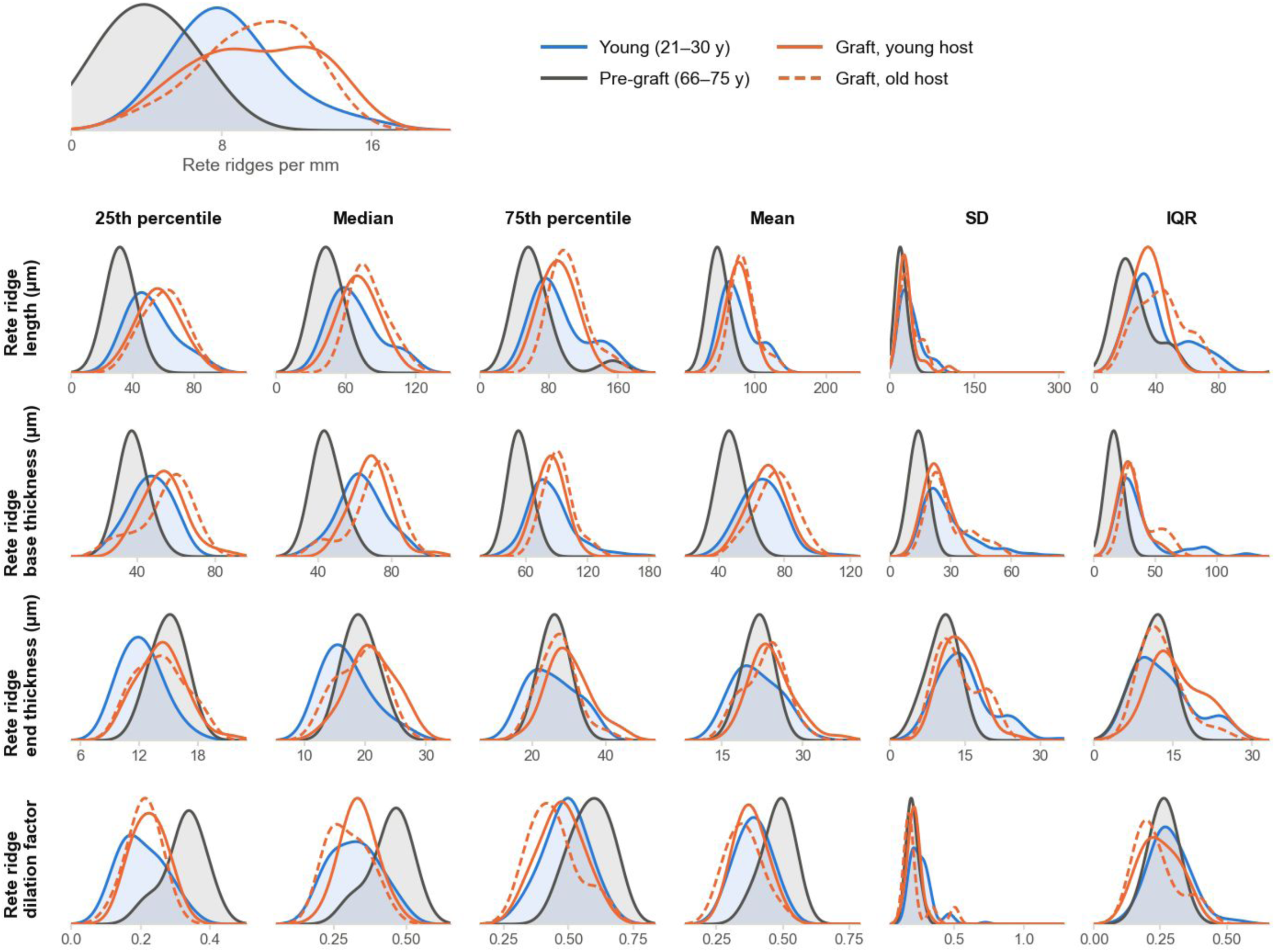
Grafts resemble never-grafted young skin across the rete ridge features that change with age. Density of per-section values, each donor weighted equally, for abdominal skin from tissue-bank donors aged 21-30 years (11 donors, 40 sections), TenBio abdominal skin from donors aged 66-75 years before grafting (7 donors, 10 sections), and the same donors’ grafts harvested from young (6 donors, 21 sections) and old hosts (5 donors, 9 sections). Top, rete ridge density. Grid, one row per rete ridge measure and one column per statistic of the ridges in each section; dilation factor is end thickness divided by base thickness.

**Supplementary Figure S11.**
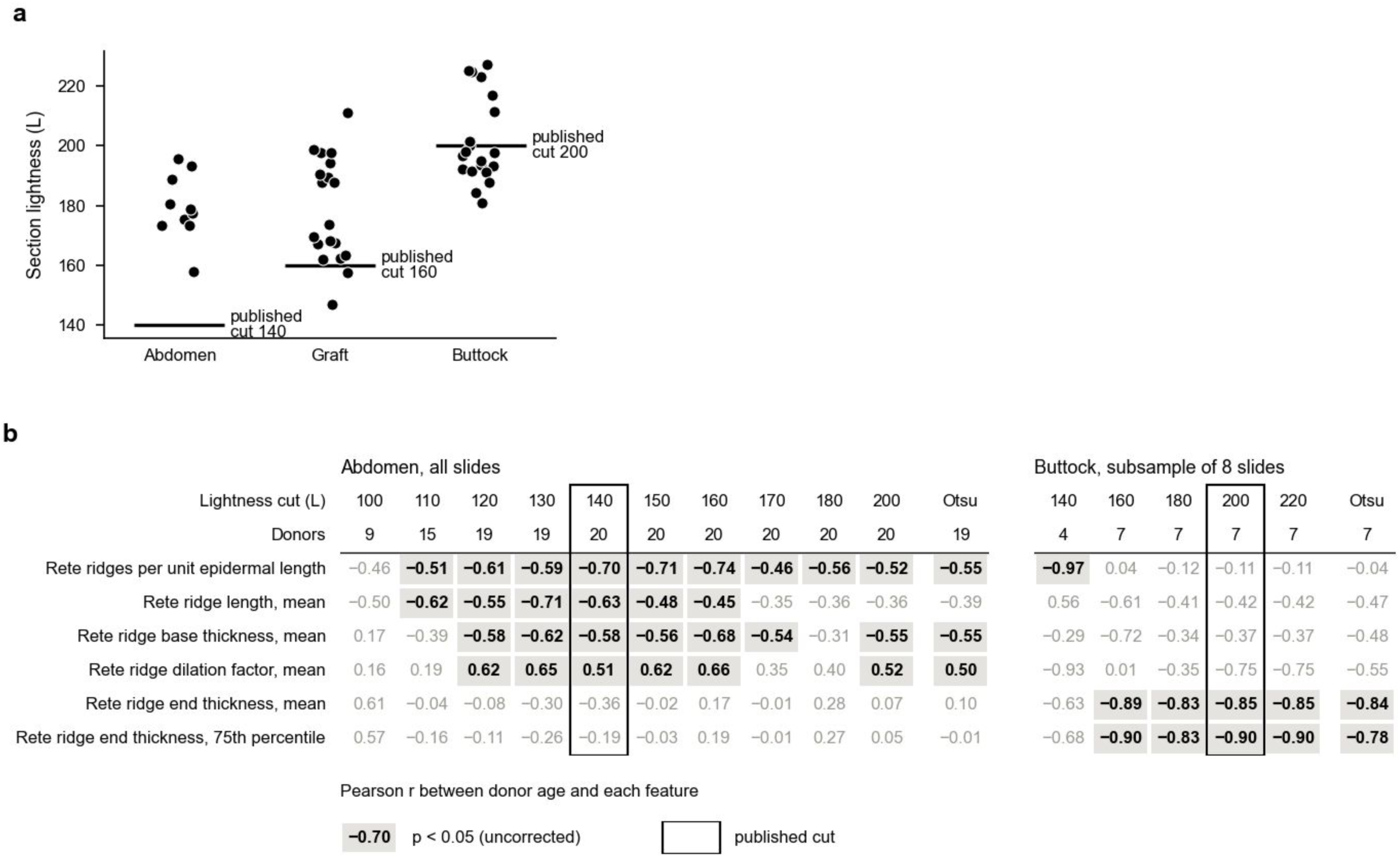
Sensitivity of the epidermal morphometrics to the lightness cut. a. Mean tissue lightness of each section by site, with each site’s published cut (L > 140 abdomen, 160 grafts, 200 buttock). b. Pearson r between donor age and each feature’s per-donor median after re-running the segmentation at ten fixed cuts (L 100 to 200) on all abdomen slides (20 donors) and five cuts (L 140 to 220) on a subsample of eight buttock slides (7 donors), and with Otsu’s method applied to each section. Shaded bold values, p < 0.05 (uncorrected); boxed column, published cut. The buttock arm shows the stability of r rather than p. Yield and feature values at every cut are in the deposited outputs.

## Notes

### Competing Interest Statement

The authors have declared no competing interest.

### Summary of Updates

Rebuilding the age prediction model without data leakage. Applying multiple testing correction to, no CD31 feature survived (smallest q = 0.087), so the vascular analysis of aging skin is now presented as a preliminary observation in the Supplement. We also explain that the vasculature could not have been measured meaningfully in grafts, since human skin is revascularized by the murine vasculature, which is why CD31 was not included in the age prediction model. Second, we corrected the statement that Ki67 did not change after grafting: a paired within-donor analysis showed that it indeed did rise in every donor with matched samples (1.80-fold into young hosts, 2.01-fold into old hosts) without reaching significance at this sample size. The revision also adds a benchmark against the two published histology-based skin clocks, a sensitivity analysis, and effect sizes with confidence intervals and q-values for every regression.

